# ACE2 utilization of HKU25 clade MERS-related coronaviruses with broad geographic distribution

**DOI:** 10.1101/2025.02.19.639017

**Authors:** Chen Liu, Young-Jun Park, Cheng-Bao Ma, Cameron Stuart, Risako Gen, Yu-Cheng Sun, Xiao Yang, Mei-Yi Lin, Qing Xiong, Jun-Yu Si, Peng Liu, David Veesler, Huan Yan

**Affiliations:** State Key Laboratory of Virology and Biosafety, College of Life Sciences, TaiKang Center for Life and Medical Sciences, Wuhan University; Wuhan, Hubei, 430072, China; Department of Biochemistry, University of Washington; Seattle, WA 98195, USA; Howard Hughes Medical Institute, University of Washington; Seattle, WA 98195, USA

**Keywords:** HKU25, MERSr-CoV, EjCoV-3, DPP4, ACE2, Receptor, Cryo-EM

## Abstract

Dipeptidyl peptidase-4 (DPP4) is a well-established receptor for several MERS-related coronaviruses (MERSr-CoVs) isolated from humans, camels, pangolins, and bats (1–6). However, the receptor usage of many genetically diverse bat MERSr-CoVs with broad geographical distributions remains poorly understood. Recent studies have identified angiotensin-converting enzyme 2 (ACE2) as an entry receptor for multiple merbecovirus clades. Here, using viral antigen and pseudovirus-based functional assays, we demonstrate that several bat merbecoviruses from the HKU25 clade previously thought to utilize DPP4 (7), employ ACE2 as their functional receptor. Cryo-electron microscopy analysis revealed that HsItaly2011 and VsCoV-a7 recognize ACE2 with a binding mode sharing similarity with that of HKU5 but involving remodeled interfaces and distinct ortholog selectivity, suggesting a common evolutionary origin of ACE2 utilization for these two clades of viruses. EjCoV-3, a strain closely related to the DPP4-using MERSr-CoV BtCoV-422, exhibited relatively broad ACE2 ortholog tropism and could utilize human ACE2 albeit suboptimally. Despite differences in entry mechanisms and spike proteolytic activation compared to MERS-CoV, these viruses remain sensitive to several broadly neutralizing antibodies and entry inhibitors. These findings redefine our understanding of the evolution of receptor usage among MERSr-CoVs and highlight the versatility of ACE2 as a functional receptor for diverse coronaviruses.

**Significance:** Recent studies unexpectedly revealed that several merbecoviruses convergently evolved ACE2 receptor usage with distinct binding modes across three continents, challenging the dogma that DPP4 is their primary receptor. Here, we demonstrate that HKU25 clade MERS-related coronaviruses broadly distributed across Eurasia utilize ACE2 as host receptor through a binding mode shared with HKU5, challenging prior findings. These findings reveal a prevalence of ACE2 usage in diverse MERS-related coronaviruses in bats and show that EjCoV-3 is preadapted to use human ACE2, suggesting a potential for spillover. Our data provide a blueprint of host receptor barrier determinants which will facilitate global surveillance and development of countermeasures against these poorly characterized merbecoviruses.

## Introduction

Middle East respiratory syndrome coronavirus (MERS-CoV) is a highly pathogenic virus with a case fatality rate of 36% (8). Since its emergence in 2012, sporadic MERS-CoV infections have been reported annually in the Middle East (9). Recently, the World Health Organization (WHO) expanded its list of prioritized coronaviruses to include the entire *Merbecovirus* subgenus, due to their epidemic and pandemic potential (10, 11). According to the International Committee on Taxonomy of Viruses (ICTV) taxonomy (August 2023)(12), the merbecovirus subgenus includes four species: *Betacoronavirus cameli* (MERSr-CoVs), *Betacoronavirus erinacei* (EriCoV), *Betacoronavirus pipistrelli* (HKU5), and *Betacoronavirus tylonycteridis* (HKU4). Although MERS-CoV is part of *Betacoronavirus cameli,* along with diverse viruses circulating in vespertilionid bats (Vespertilionidae), there is a phylogenetic gap connecting these merbecoviruses (13, 14). The closest known relative of human and camel MERS-CoV is NeoCoV, which was discovered in *Neoromicia capensis* (Cape serotine bat) in Africa and only shares 85.5% whole genome nucleotide sequence identity with MERS-CoV and exhibits significant divergence in the Spike (S) glycoprotein S_1_ subunit (15–17).

DPP4 was first identified as the entry receptor for MERS-CoV in 2013 and was later shown to also mediate entry of HKU4-related viruses (*Betacoronavirus tylonycteridis*), which includes strains from Tylonycteris bats and pangolins (1–5). While HKU4 and a few bat MERSr-CoVs, such as BtCoV-422 (6), share similar RBD features with human/camel MERS-CoV, many other MERSr-CoVs exhibit highly divergent receptor-binding domain (RBD) sequences, suggesting the use of alternative receptors (18). Indeed, the extraordinary genetic diversity observed in merbecovirus RBDs emphasizes the challenges associated with predicting zoonotic risks of these viruses (14, 18, 19). As a result, we classified the phylogenetic diversity of merbecovirus RBDs into six distinct clades to provide a framework to understand receptor usage and support vaccine design and pandemic preparedness efforts (20).

We and others recently revealed that merbecoviruses from the NeoCoV, MOW15-22, and HKU5 clades, comprising viruses found on three continents, have independently evolved the ability to utilize ACE2 as a receptor using entirely distinct binding modes (17, 19–23). The receptor switch history of merbecovirus remains unclear but recombination appears to play a crucial role in these events (14, 17, 24, 25). Therefore, merbecovirus receptor usage can markedly deviate from the taxonomy of viral species based on the conservation of five concatenated replicase domains in ORF1ab (26, 27).

Although bat MERSr-CoV HKU25 has been proposed to use DPP4 for entry, the supporting data is not strong, and structural evidence supporting this claim is lacking(7). Consequently, there is uncertainty as to the nature of the receptor used for cell entry by merbecoviruses from the HKU25 clade, including viruses discovered in Italy (28), Switzerland (29), China (6, 7, 30, 31), and Japan(32), limiting our ability to predict the spillover potential of these important pathogens. Here, we hypothesized, that members of the HKU25 clade of coronaviruses utilize ACE2 rather than DPP4 as their receptor based on phylogenetic relatedness to HKU5. Screening an ACE2 ortholog library revealed that most, but probably not all, HKU25 clade coronaviruses can engage ACE2 from several bat species and a subset of non-bat mammals, particularly those from the Artiodactyl and Rodent orders. EjCoV-3, a strain closely related to the DPP4-using MERSr-CoV BtCoV-422 at the whole genome level, demonstrated broad ACE2 ortholog tropism and a weak ability to utilize human ACE2 (hACE2). Cryo-electron microscopy analysis of the ACE2-bound HsItaly2011 and VsCoV-a7 RBDs showed that these viruses engage ACE2 with a binding pose reminiscent of that observed for HKU5, but involving remodeled interfaces and distinct ortholog selectivity, suggesting a common evolutionary origin of ACE2 utilization for these two clades of viruses (20).

## Results

### Prediction of ACE2 utilization by HKU25 clade coronaviruses

To investigate receptor usage among diverse merbecoviruses, we retrieved publicly available β-coronavirus S sequences from the National Center for Biotechnology Information (NCBI) database. Phylogenetic analysis based on amino acid sequences identified 1,117 S sequences classified as merbecoviruses. After removing redundant sequences with identical amino acid compositions and over-sampled human MERS-CoV strains, we selected 152 S sequences (***SI Appendix,* Dataset S1**) for multiple sequence alignment and phylogenetic tree construction to identify representative strains (***SI Appendix,* Fig. S1*A***). Further phylogenetic analyses of S (***SI Appendix,* Fig. S1*B***) and receptor-binding domain (RBD) sequences (**Fig. 1*A***) were conducted on representative strains spanning four species, with a focus on viruses without confirmed receptors. These included the bat coronavirus NsGHA2010 (33), hedgehog coronaviruses (EriCoVs) (34–36), and 15 non-redundant bat coronaviruses classified as members of the HKU25 clade (6, 7, 31). Comparative analysis of trees based on whole-genome nucleotide sequences and S/RBD amino acid sequences revealed phylogenetic incongruencies (**Fig. 1*B***). For example, the three geographically separated MERSr-CoV strains EjCoV-3 (32), BtCoV-422 (6, 7), and VmSL2020 (29), which exhibit significant divergence in their S/RBD region, clustered together and share 84.5∼89.4% genome-wide nucleotide sequence identity. Analysis of amino acid sequences from five concatenated domains in the replicase region (3CLpro, NiRAN, RdRp, ZBD, and HEL1) within ORF1ab confirmed that all HKU25 clade coronaviruses are classified as MERSr-CoVs (>92.4% identity compared to MERS-CoV) (26, 27) (**Fig. 1*B***). Phylogenetic analysis of RBD sequences revealed a close relationship between HKU5- and HKU25 clade coronaviruses, suggesting that members of the HKU25 clade of coronaviruses may also utilize angiotensin-converting enzyme 2 (ACE2) as receptor, similar to HKU5 (20, 22, 23). Whereas HKU5 was predominantly sampled in *Pipistrellus abramus* (*P.abr*) bats in Southeast China, HKU25 clade coronaviruses have been identified in a wide range of vespertilionid bat species across Eurasia. These include: VmSL2020 and VmSL2021 from *Vespertilio murinus* (*V.mur*) in Switzerland (29); PaGB01 from *Plecotus auritus* (*P.aur*) in the United Kingdom (37), HsItaly2011 from *Hypsugo savii* (*H.sav*), and PkItaly2011 from *Pipistrellus kuhlii* (*P.kuh*) in Italy (28), SC013 from *Vespertilio superans* (*V.sup*) (30), GD2016-Q249 from *Pipistrellus abramus* (*P.abr*) (31); HKU25 strains from *Hypsugo pulveratus* (*H.pul*) (6) in China; VsCoV-1, VsCoV-kj15, VsCoV-a7 from *Vespertilio sinensis* (*V.sin*, same species as *Vespertilio superans*) and EjCoV-3 from *Eptesicus japonensis* (*E.jap*) in Japan (32) (**Fig. 1*C***).

**Fig. 1.**
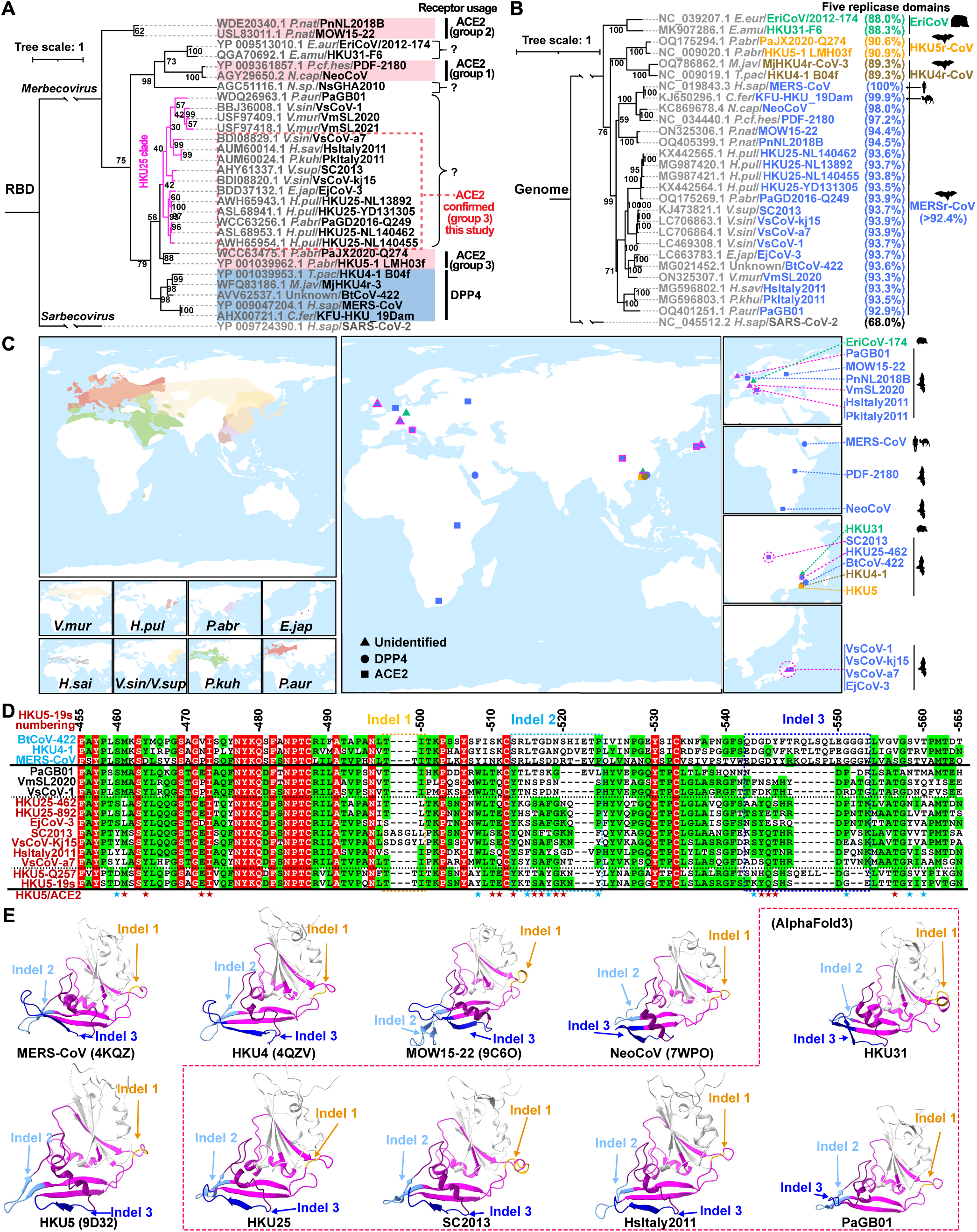
Genetic features and geographic distribution of HKU25 clade MERSr-CoVs. (***A-B***) Maximum-likelihood phylogenetic trees of representative merbecoviruses, generated using IQ-tree2. Trees are based on amino acid sequences of the RBD (*A*) or genomic nucleotide sequences (*B*), with SARS-CoV-2 as the outgroup. Information on receptor usage, binding mode, host, and amino acid sequence identities of five replicase domains (3CLpro, NiRAN, RdRp, ZBD, and HEL1) for coronavirus taxonomy (MERSr-CoV defined as diverged by less than 7.6% identity to MERS-CoV, NC_019843.3) are annotated. The known ACE2-using viruses were classified according to the three distinct binding modes identified in the NeoCoV- (group 1), MOW15-22- (group 2), and HKU5-related (group 3) clades (27). The scale bars denote genetic distance (1 substitution per nucleotide/amino acid position). (***C***) Geographic distributions of bat hosts (left) and sampling locations of merbecoviruses with annotated receptor usage (right). Data from the IUCN (International Union for Conservation of Nature) Red List were visualized using GeoScene Pro. Squares: ACE2-using; Circles: DPP4-using; Triangles: receptor-unidentified. Color coding is the same as panel 1B. HKU25 clade strains were outlined in magenta. Host abbreviations: *V.mur*/*V.sup* (*Vespertilio murinus*/*V.superans*), *H.pul* (*Hypsugo pulveratus*), *E.jap* (*Eptesicus japonensis*), *H.sai* (*Hypsugo savii*), *V.sin* (*Vespertilio sinensis*), *P.kuh* (*Pipistrellus kuhlii*), *P.aur* (*Plecotus auritus*), *P.abr* (*Pipistrellus abramus*). (***D***) RBM sequence alignment of the indicated merbecoviruses with manual adjustment to optimize indel positioning. Fully and partially conserved residues were indicated as red and green backgrounds, respectively. Dashed boxes highlight indels. Residues involved in HKU5-ACE2 interactions are marked with stars; positions that are conserved or non-conserved compared to HKU25 clade viruses are colored in red and blue, respectively. HKU5-19s residue numbering is shown. (***E***) Experimentally determined structures or AlphaFold3-predicted RBD structures of representative merbecoviruses. The putative RBMs are indicated in magenta and three featured indels described in panel D are labeled in orange (indel 1), light blue (indel 2), and dark blue (indel 3), respectively. Sequences between indel 2 and indel 3 are labeled in purple to facilitate observation.

Pairwise amino acid sequence analysis showed that S glycoproteins from HKU25 clade coronaviruses share 65-68% identities with MERS-CoV S, 67-70% identities with HKU4-1 S, and 69-73% identities with HKU5-1 S, respectively. Furthermore, HKU25 clade RBDs share 48-54% identity with HKU4-1 and 62-73% with HKU5-1, but markedly lower homology (32-36%) with NeoCoV and MOW15-22, suggesting distinct receptor recognition modes(17, 19) (**Fig. S1C**). HKU25 clade RBDs harbor insertions-deletions (indels) similar to that found in HKU5 (e.g. two indels at HKU5 S residues 513-522 and 543-552, respectively), but distinct from that of MERS-CoV, HKU4-1, BtCoV-422, NeoCoV, MOW15-22, and HKU31 (***SI Appendix,* Fig. S1D**). Additional insertions-deletions (indels) can be found in PaGB01, SC2013, VsCoV-kj15, and other viruses. Up to 15 out of 24 ACE2-interacting HKU5-19s residues are conserved in HKU25 clade RBDs, suggesting a potentially shared receptor usage (**Fig. 1*D***). Simplot analysis comparing several viral genome sequences with EjCoV-3 reveals high similarity to BtCoV-422 and VmSL2020 with marked divergence in the S_1_ region. Accordingly, the EjCoV-3 RBD is more closely related to the ACE2-using HKU5-1 RBD than to the DPP4-utilizing BtCoV-422 RBD (1), suggesting possible recombination events among ancestral strains (***SI Appendix,* Fig. S1*E***). Furthermore, AlphaFold3-predicted structures of HKU25-related RBDs highlight their similarity to HKU5-1 in terms of overall RBM architecture, especially the RBM indel 2 and 3 located at the tip (20), except for PaGB01 which harbors a short indel 3 (**Fig. 1*E***).

Overall, these findings suggest that members of the HKU25 clade of MERSr-CoVs may use ACE2 as host receptor through a binding mode similar to that of HKU5 (20), setting them apart from DPP4-using MERS-CoV and HKU4 clade coronaviruses or other ACE2-using MERSr-CoVs.

### Multi-species ACE2 tropism of HKU25 clade coronaviruses

To investigate the receptor usage of HKU25 clade coronaviruses, we first tested binding of RBD-human IgG Fc (RBD-hFc) fusion constructs from eleven HKU25 clade strains to ACE2 and DPP4 orthologs from *P.aur, P.kuh, V.mur,* and *P.abr*, which are reported host species of HKU25 clade coronaviruses. None of these ACE2 or DPP4 orthologs supported RBD-hFc binding of virus strains (PaJX2020-Q274, PkItaly2011, VmSL2020, VmSL2021, and PaGB01) identified in the corresponding host species. However, *P.aur* ACE2, but not DPP4, unexpectedly bound RBD-hFc constructs from six HKU25 clade strains efficiently (***SI Appendix,* Fig. S2**).

To further explore the multi-species ACE2 tropism of HKU25 clade coronaviruses, we subsequently assessed RBD-hFc binding and VSV pseudovirus entry of eight representative strains using a well-established ACE2 library comprising 113 ACE2 orthologs from 59 bats and 54 non-bat mammalian species with validated expression(21) (**Fig. 2*A* and *B*, and *SI Appendix,* Fig. S3**). We found that HKU25, EjCoV-3, SC2013, HsItaly2011, and VsCoV-a7 can efficiently bind to and utilize multiple ACE2 orthologs from bats and non-bat mammalian species with an overall preference for *Murina feae* (*M.fea*), *Eptesicus fuscus* (*E.fus*), and *P.aur* bat ACE2 orthologs along with several Artiodactyl and Rodent ACE2s. Although the EjCoV-3 RBD-hFc only weakly bound to select ACE2 orthologs, EjCoV-3 S VSV pseudovirus exhibited a broad ACE2 tropism across diverse mammalian orders and was the only HKU25 clade coronavirus tested capable of utilizing hACE2 for cell entry. In contrast, we did not detect meaningful binding or pseudovirus entry for any ACE2 ortholog tested with VmSL2020, VsCoV-1, or PaGB01 (**Fig. 2*A* and *B***).

**Fig. 2.**
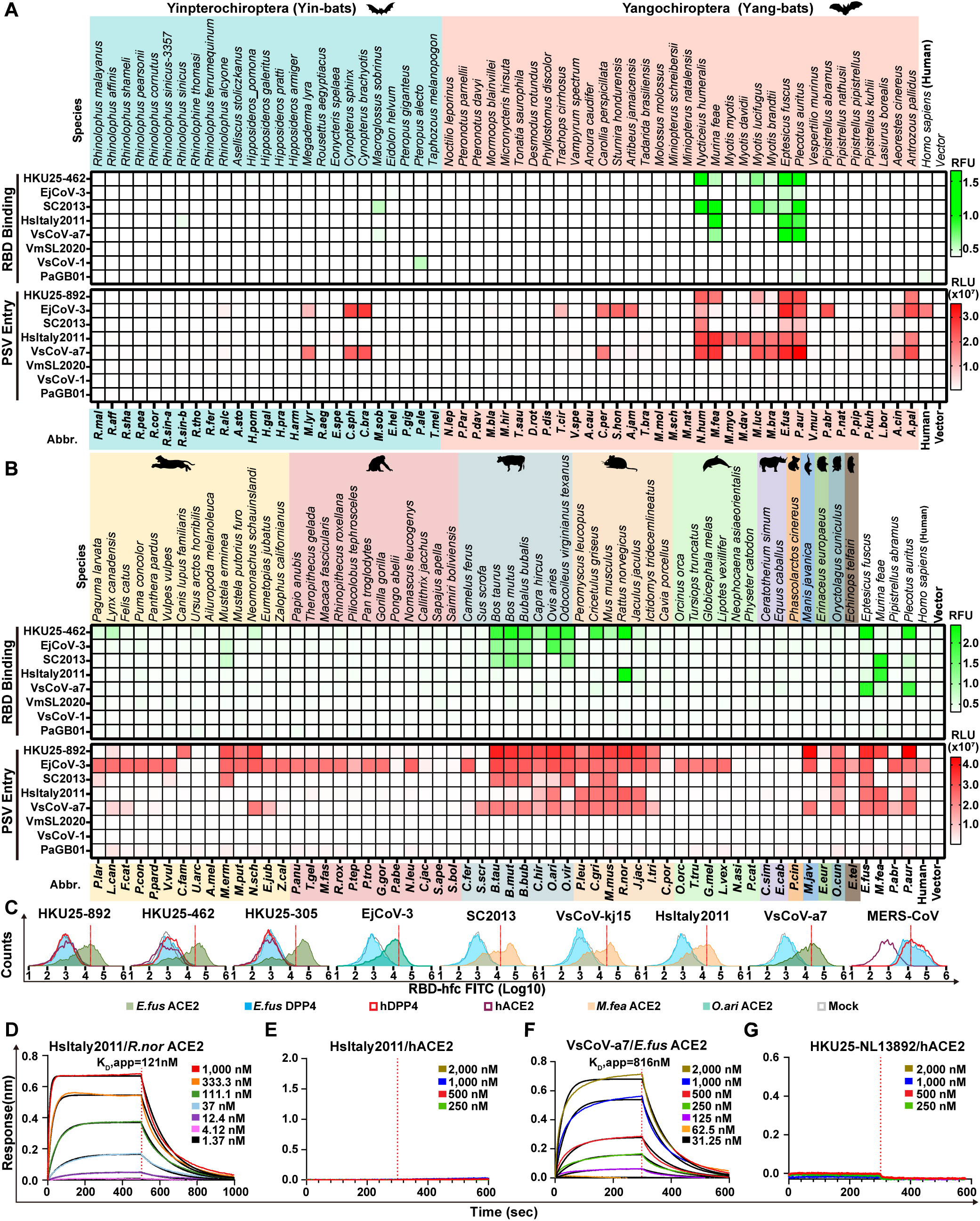
ACE2 ortholog utilization of HKU25 clade MERSr-CoVs. (***A* and *B***) Heat map representing the magnitude of HKU25 clade RBD-hFc binding to (green) and VSV pseudovirus (PSV) entry into (red) HEK293T cells transiently expressing bat (A) or non-bat (***B***) mammalian ACE2 orthologs. Mammalian orders are color-coded: Carnivora, Primates, Artiodactyla, Rodentia, Cetacea, Perissodactyla, Diprotodontia, Pholidota, Erinaceomorpha, Lagomorpha, Chiroptera. Data represent mean values (n = 3 biological replicates). PSVs were pretreatment with 100 μg/ml TPCK-treated trypsin (Try). (***C***) Flow cytometry analysis of binding of HKU25 clade RBDs to HEK293T transiently expressing the indicated ACE2 or DPP4 orthologs. Grey: vector control. Dashed lines: background threshold. Data are means of three technical repeats from three tubes of cells. (***D-G***) BLI analysis of binding kinetics of dimeric ACE2 ectodomains (*R.nor* ACE2 in panel *D*, *E.fus* ACE2 in panel *F*, and hACE2 in panel *E*/*G*) to immobilized RBD-hFc of indicated strains. Analysis was conducted using curve-fitting kinetic with global fitting (1:1 binding model).

Concurring with the above data, flow cytometry analysis further showed that the RBD-hFc of HKU25 clade coronaviruses bound to ACE2 orthologs from E.fus and M.fea, but not to human or *E.fus* DPP4, contradicting a previous report which proposed that HKU25 can bind and utilize hDPP4 (7, 21) (**Fig. 2*C***). Using biolayer interferometry (BLI), we found that the soluble dimeric *R.nor* ACE2 bound to the immobilized HsItaly2011 RBD with apparent affinity (K_D_,app) of 121 nM and that the dimeric *E.fus* ACE2 ectodomain bound to the immobilized VsCoV-a7 RBD with a K_D_,app of 816 nM. We could not detect binding of dimeric hACE2 ectodomain to either RBDs (**Fig. 2*D-G***). Together, these results demonstrate that a subgroup of HKU25 clade coronaviruses utilize ACE2 as entry receptor while largely excluding the role of DPP4 in cell entry.

### Molecular basis of HsItaly2011 and VsCoV-a7 utilization of ACE2

To understand the molecular basis of HKU25 clade coronavirus engagement of ACE2 host receptors, we characterized the *E.fus* (Bat) ACE2-bound VsCoV-a7 and *R.nor* (Rodent) ACE2-bound HsItaly2011 RBDs complexes using single particle cryoEM (**Fig. 3*A*, *SI Appendix,* Fig. S4-S5 and Table S1**). The use of natively dimeric ACE2 ectodomain constructs enabled leveraging the applied C2 symmetry to perform symmetry expansion yielding reconstructions at 2.5 Å resolution.

**Fig. 3.**
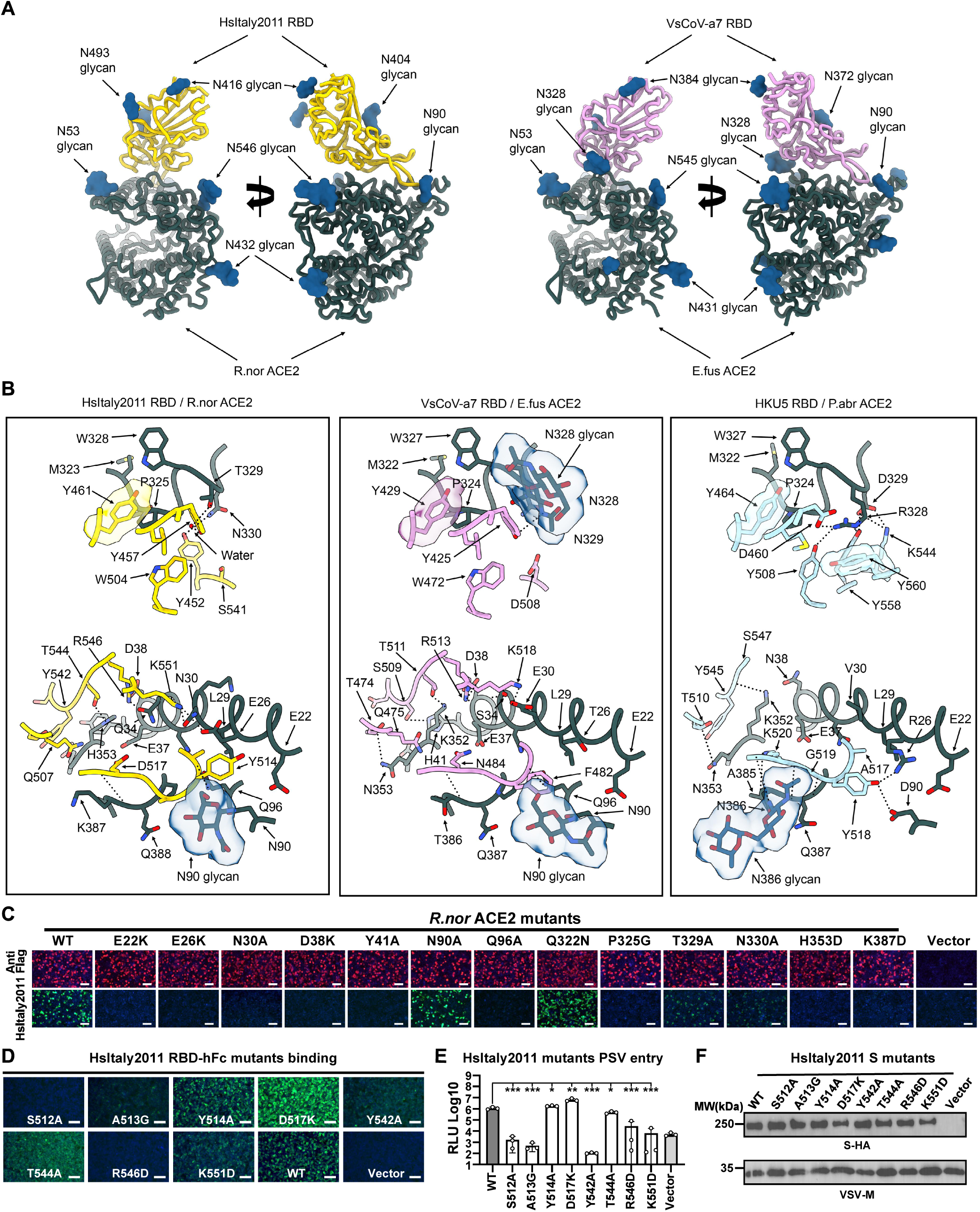
Structural basis for HsItaly2011 and VsCoV-a7 RBD interaction with bat or rat (*R.nor*) ACE2 orthologs. (***A***) Ribbon diagrams in two orthogonal orientations of the cryo-EM structures of the *R.nor* ACE2 peptidase domain (green) bound to HsItaly2011 RBD (gold) and *E.fus* ACE2 peptidase domain (green) bound to VsCoV-a7 RBD (plum). (***B***) Zoomed-in views and comparisons of the interface key interactions of the HsItaly2011 RBD/*R.nor* ACE2, VsCoV-a7 RBD/*E.fus* ACE2 and HKU5 RBD/*P.abr* ACE2 (PDB ID: 9D32). HKU5 RBD and *P.abr* ACE2 peptidase domain are colored in light blue and green, respectively. Selected interface interactions are shown as black dotted lines. (***C***) Analysis of HsItaly2011 RBD-hFc binding to membrane-anchored wildtype and mutants *R.nor* ACE2 transiently transfected in HEK293T cells analyzed by immunofluorescence. (***D*** and ***E***) RBD-hFc binding (*D*) and pseudovirus (pretreatment with 100 μg/ml TPCK-treated trypsin) entry (*E*) efficiencies of HsItaly2011 S mutants in HEK293T cells transiently expressing *R.nor* ACE2. (***F***) VSV packaging efficiencies of HsItaly2011 S mutants. VSV-M was used as a loading control. Unpaired two-tailed t-tests for *E*, data are presented as means ± SD for for n = 3 biological replicates. \**P* < 0.05,\*\**P* < 0.01, \*\*\**P* < 0.005. Scale bars in *C* and *D*: 100 μm. RLU: relative light unit.

Strikingly, the VsCoV-a7 and HsItaly2011 RBDs engage the ACE2 peptidase domain with comparable binding poses to that recently described for the ACE2-bound HKU5 RBD complex with which they can be superimposed with r.m.s.d. values of 0.9 and 1.0 Å over 776 and 774 aligned Cα positions. The interfaces of VsCoV-a7 RBD - *E.fus* ACE2 complex and HsItaly2011 RBD - *R.nor* ACE2 complexes bury an average surface of 875 Å^2^ and 1018 Å^2^, respectively, as compared to 950 Å^2^ for the HKU5 RBD - *P.abr* ACE2 complex (20).

*E.fus* ACE2 and *R.nor* ACE2 respectively interact with the VsCoV-a7 RBD and the HsItaly2011 RBD through both shared interactions and contacts specific to each complex involving the molecular determinants of receptor species tropism previously identified for HKU5. For instance, P324*_E.fus_*_ACE2_, P325*_R.nor_*_ACE2_, and P324*_P.abr_*_ACE2_ insert in a comparable crevice at the surface of each RBM through Y429_VsCoV-a7_, Y461_HsItaly2011_, and Y464_HKU5-19s_. Conversely, neighboring interactions are profoundly remodeled relative to HKU5, as exemplified by N328*_E.fus_*_ACE2_ harboring an N-linked oligosaccharide, which is directly interacting with Y425_VsCoV-a7_, or T329*_R.nor_*_ACE2_ which is hydrogen-bonded to the equivalent Y457_HsItaly2011_ residue through a water molecule **(Fig. 3B)**. The N353*_E.fus_*_ACE2_ side chain is hydrogen-bonded to the T474_VsCoV-a7_ backbone carbonyl (similar to N353*_Pabr_*_ACE2_ and T510_HKU5_) whereas G354*_R.nor_*_ACE2_ cannot form similar interactions with the HsItaly2011 RBD. The VsCoV-a7 and HsItaly2011 RBDs respectively accommodate an ACE2 glycan at position N90*_E.fus_*_ACE2_ and N90*_R.nor_*_ACE2_ through F482_VsCoV-a7_ and Y514_HsItaly2011_, setting them apart from the *P.abr* ACE2 - HKU5 interface (**Fig. 3*B***).

To functionally validate the contribution of the identified interacting residues in receptor recognition, we examined the influence on HsItaly2011 RBD-hFc binding of *R.nor* ACE2 substitutions at key positions (**Fig. 3*C***). Most point mutations evaluated reduced HsItaly2011 RBD-hFc binding with the exception of the N90A*_R.nor_*_ACE2_ glycan knock-out and Q322N*_R.nor_*_ACE2_ glycan knock-in mutations, suggesting a minor role of these glycans in modulating HsItaly2011 receptor engagement (**Fig. 3*C***). Consistent with the structural data, several alanine/glycine substitutions or charge reversals in the HsItaly2011 RBM dampened RBD-hFc binding and pseudovirus entry efficiency except for the Y514A, D517K, and T544A mutations, probably due to the remodeling of the contacts (**Fig. 3*D-F***).

### Critical residues and glycans for ACE2 tropism determination

To investigate the host species tropism determinants of HKU25 clade coronaviruses, we engineered chimeric ACE2 constructs by swapping sequence regions between functional orthologs from *P.aur*, *M.erm*, and the non-functional hACE2. Our structural analysis revealed that direct virus-interacting amino acids are located between residues 1-100 and 301-400. We therefore generated chimeras by swapping three consecutive regions: residues 1-100, 101-300, and 301-400. Swaps in residues 101-300 had minimal impact on receptor functionality compared to wild-type (WT) controls whereas substitutions in residues 1-100 or 301-400 altered receptor functionality **(Fig. 4*A*)**. For example, chimeras containing hACE2 residues 1-100 lost their RBD-binding ability, indicating the crucial role of these residues in receptor recognition. We further set out to determine the molecular determinants restricting the functionality of hACE2. Replacing residues 1-100 in hACE2 with sequences from *P.aur* or *M.erm* ACE2 restored binding to several HKU25 clade RBDs. Swaps in residues 301-400 caused milder phenotypic changes (**Fig. 4*A***). However, swaps of residues 1-50 (relative to *P.aur* ACE2) or substituting five key residues (relative to *M.feas* ACE2) failed to rescue hACE2 functionality (**Figs. 4*B* and *SI Appendix,* S6*A* and *B***). Through testing several hACE2 mutations within residues 301-400, we identified that the Q325P mutation promoted detectable HKU25-NL140462 RBD binding which was further enhanced by the additional E329R/N330D substitutions, more closely matching the *P.abr* ACE2 residues (HKU5 receptor, **Fig. 4*B***). Moreover, *M.erm* ACE2 residue 354, a critical determinant of HKU5 receptor species tropism (20), also influenced EjCoV-3 and VsCoV-a7 RBD binding efficiency, underscoring shared molecular determinants between HKU25 and HKU5 clades (***SI Appendix,* Fig. S6*C***).

**Fig. 4.**
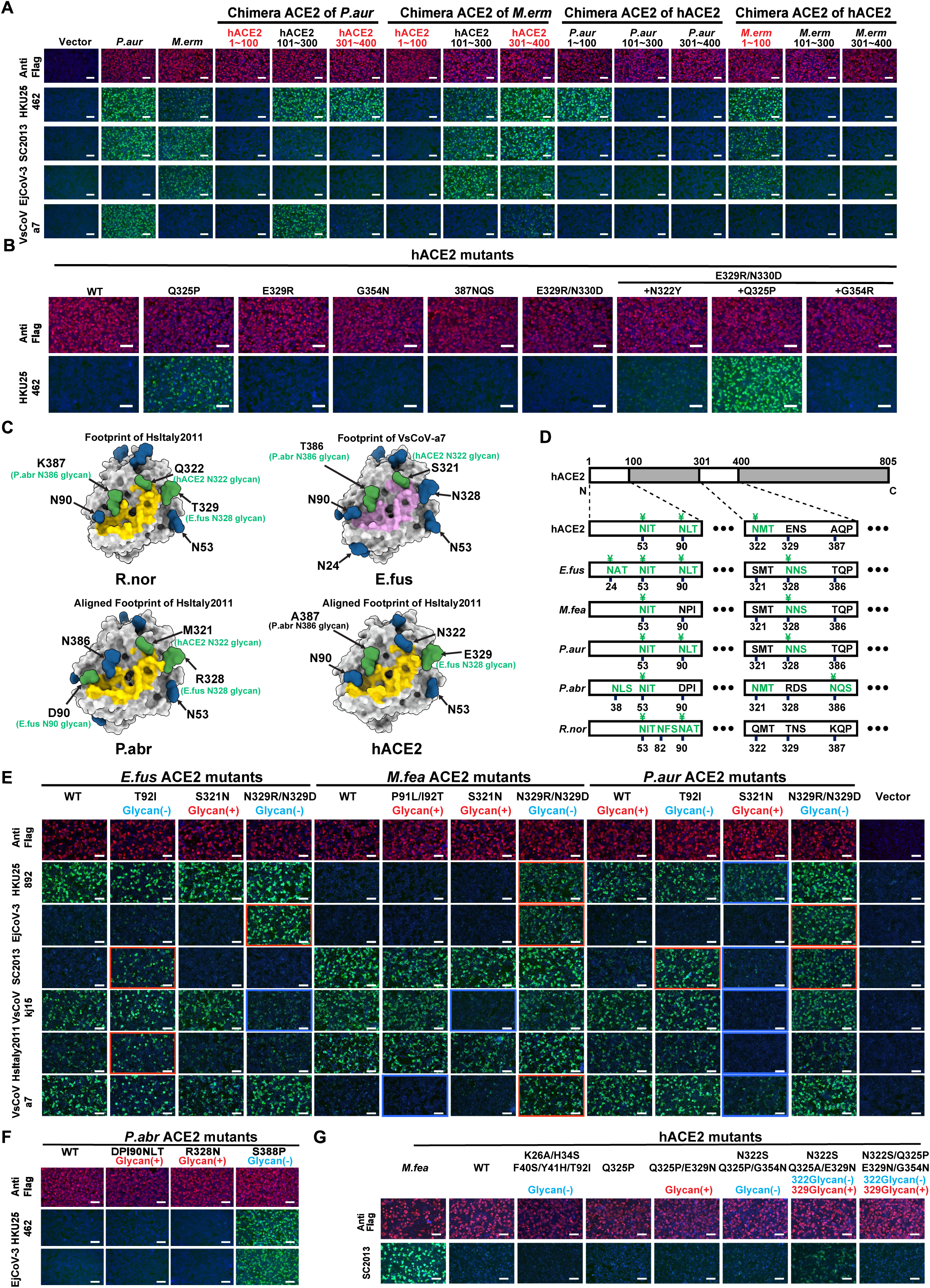
ACE2 tropism determinants for HKU25 clade coronavirus. (***A***) Immunofluorescence analysis of RBD-hFc binding to HEK293T cells transiently expressing ACE2 chimeras (swaps between hACE2/*P.aur* ACE2 or hACE2/*M.erm* ACE2. ACE2 expression were validated by detecting the C-terminal fused FLAG tags. (***B***) HKU25-NL140462 RBD-hFc binding to hACE2 mutants with equivalent residues in *P.abr* ACE2. (***C***) N-glycans proximal to or within the HKU25 clade RBD-ACE2 interface (residues 1–100, 301–400). HsItaly2011 (yellow) and VsCoV-a7 (pink) RBD footprints are mapped onto ACE2 orthologs (gray surface). Glycans actually present on the surface of indicated WT ACE2 orthologs or predicted glycans through glycan-knock in mutations (based on hACE2, PDB 6M0J) are rendered in blue and green, respectively. (***D***) Glycosylation sequons (green) at positions 53, 90, 322, 329, and 387 (hACE2 numbering). Cryo-EM confirmed glycans are marked with ¥. Please note although several glycosylation sequons are present, no glycan is present in these sites according to the cryo-EM map. (***E-G***) RBD-hFc binding assay evaluating the impact of N-Glycan mutations on *E.fus*/*M.fea*/*P.aur* ACE2 (*E*), *P.abr* ACE2 (*F*) or hACE2 (*G*) orthologs. Red/blue dashed outlines: enhanced/reduced binding. Scale bars: 100 µm.

Prior studies have highlighted the key role that some ACE2 N-linked glycans located near RBD-interacting interfaces can play in modulating receptor recognition through binding enhancement or steric restriction (17, 19, 20). To assess the functional impact of ACE2 glycans on HKU25 clade receptor utilization, we mutated each of the four glycosylation sites (hACE2 numbering positions 90, 322, 329, and 387) near or within the interaction interface (**Fig. 4*C* and *D*)**. HKU25 clade RBD binding assays showed that removing glycans at these sites can enhance binding to varying degrees in several mutants, while glycan knock-in abolished binding in several mutants (**Fig. 4*E***). Furthermore, the glycan knockout *P.abr* ACE2 S388P and the hACE2 N322S/Q325A/E329N mutations promoted detectable HKU25-NL140462/EjCoV-3 and SC2013 RBD binding, respectively (**Fig. 4*F* and *G***). These results suggest that glycans primarily act as host-tropism barriers for HKU25 clade coronaviruses, as opposed to promoting binding as is the case for the NeoCoV-*P.pip* ACE2 interaction (17).

In summary, ACE2 ortholog specificity and host range of HKU25 clade coronaviruses are governed by key critical interface residues and modulated by glycan shields.

### Characterization of S-mediated entry of HKU25 clade coronaviruses

A notable difference between HKU25-related coronaviruses and HKU5 or MERS-CoV is the absence of a polybasic (furin) cleavage site at the S_1_/S_2_ junction, a feature critical for proteolytic processing during viral biogenesis (38, 39) (**Fig. 5*A***). Accordingly, pseudotyped particles carrying HKU25 clade S glycoproteins were uncleaved when produced in HEK293T cells, with the exception of EjCoV-3 S exhibiting minimal cleavage (**Fig. 5*B***). We found that HKU25 clade S glycoproteins promoted cell-cell fusion in Caco-2 cells expressing functional ACE2 orthologs from several mammalian species, but not with hACE2, in a trypsin-dependent manner, highlighting a requirement for exogenous protease priming under the tested conditions (**Fig. 5*C***).

**Fig. 5.**
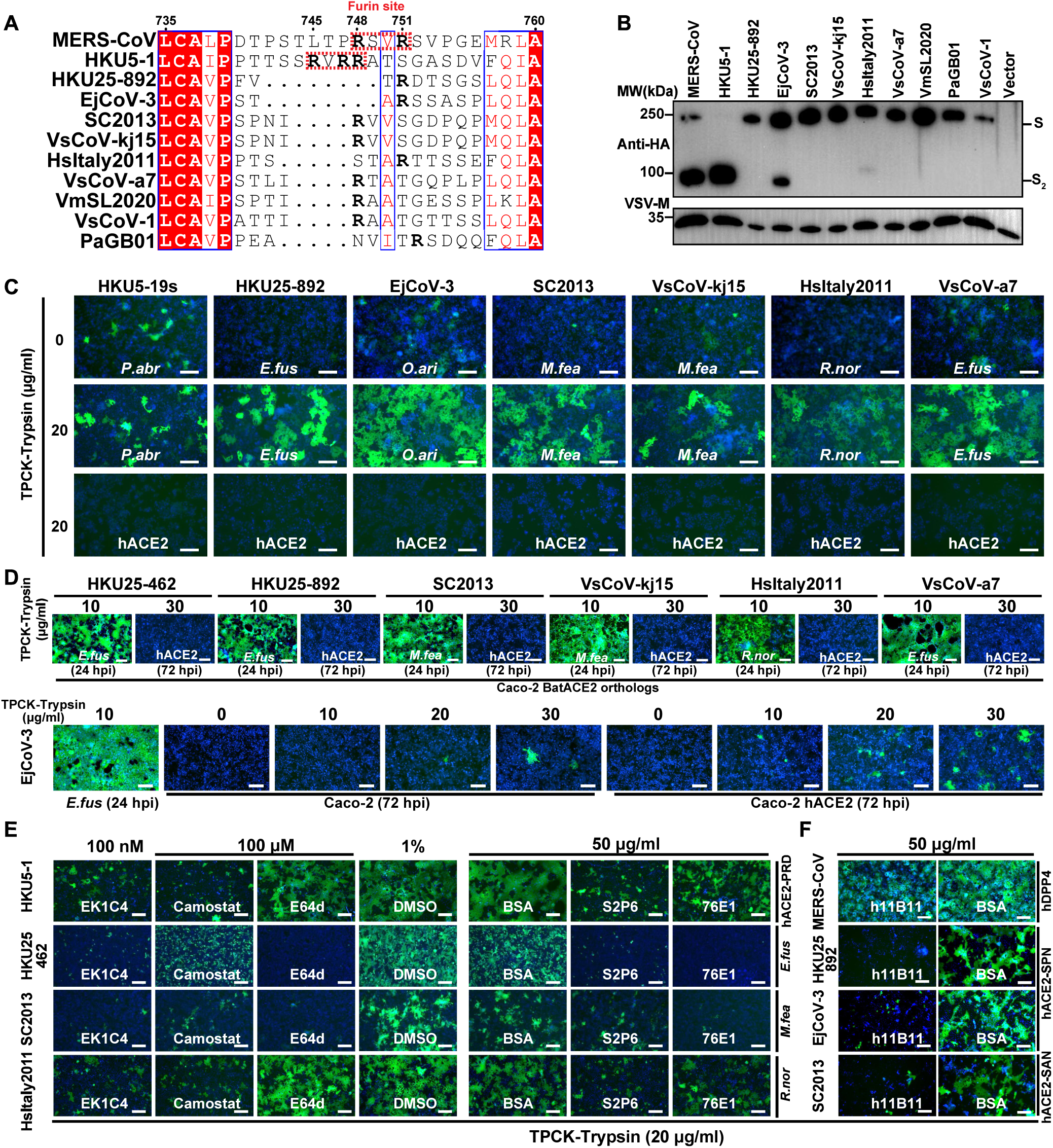
The Characterization and inhibition of the ACE2-mediated entry of rcVSV pseudotypes with HKU25 clade S glycoproteins. (***A***) S_1_/S_2_ junction sequence alignment of HKU25 clade S glycoproteins with MERS-CoV residue numbering. The arginines (R) were highlighted in bold fronts. Furin cleavage sites are highlighted in red dashed boxes. (***B***) S glycoprotein incorporation into VSV pseudoviral particles by detecting the C-terminal fused HA tags. VSV-M was used as a loading control. (***C***) Cell-cell fusion mediated by HKU25 clade coronaviruses S in Caco-2 cells stably expressing indicated ACE2 orthologs with the treatment of TPCK-treated trypsin. (***D***) Propagation of rcVSV-HKU25r-S in Caco-2 cells or Caco-2 hACE2 cells in the presence of the indicated concentrations of TPCK-treated trypsin. hpi: hours post-infection. (***E* and *F***) Inhibition of rcVSV-HKU25r-S propagation by small molecular inhibitors, S_2_ antibodies (*E*) or hACE2-targeting antibodies h11B11 (*F*) in Caco-2 cells stably expressing indicated hACE2 mutants. BSA: Bovine serum albumin, 50 μg/ml. PRD mutation: Q325P/E329R/N330D; SPN mutation: N322S/Q325P/G354N; SAN mutation: N322S/Q325A/E329N. Scale bars: 200 μm.

To evaluate S-mediated viral propagation in human cells, we used a replication-competent VSV-CoV-S (rcVSV-S) pseudotyping system in Caco-2 cells expressing human or bat ACE2 orthologs. Seven rcVSV-HKU25r-S viruses were rescued and amplified efficiently in Caco-2 cells stably expressing ACE2 orthologs from *E.fus*, *M.fea*, or *R.nor* (**Fig. 5*D***). Moreover, only EjCoV-3 exhibited detectable (weak) propagation in Caco-2 cells endogenously expressing hACE2 or overexpressing hACE2 at 72 hours post-infection (hpi), consistent with the results from single-round VSV-S pseudovirus entry assays (**Fig. 2*A* and *B***).

To delineate entry pathways and therapeutic targets, we tested rcVSV-HKU25r-S amplification in the presence of inhibitors or neutralizing antibodies targeting distinct steps of the coronavirus entry pathway. Broadly neutralizing antibodies S2P6 (40) and 76E1 (41) against the S_2_ subunit, and the OC43 HR2-derived EK1C4 lipopeptide (42, 43) effectively suppressed propagation of rcVSV-HsItaly2011-S, rcVSV-SC2013-S, and rcVSV-HKU25 NL140462-S. However, sensitivity to the cathepsin B/L inhibitor E64d and the TMPRSS2 inhibitor Camostat (44, 45), varied across rcVSV-S pseudoviruses, suggesting distinct host protease preference and entry pathways among HKU25 clade coronaviruses (**Fig. 5*E***). Additionally, the hACE2-targeting monoclonal antibody h11B11 (46, 47) neutralized viruses bearing S glycoproteins from HKU25-NL13892, EjCoV-3, and SC2013, but not MERS-CoV, in Caco-2 cells or Caco-2 cells expressing the indicated hACE2 mutants (**Fig. 5*F***). These findings underscore the potential of broad-spectrum entry inhibitors and antibodies as countermeasures against zoonotic spillovers of ACE2-using HKU25 clade MERSr-CoVs.

## Discussion

DPP4 had been established as a canonical receptor for *Merbecovirus* subgenus members since its identification as entry receptor for MERS-CoV. While this likely applies to the entire HKU4 clade, only a limited number of MERS-related coronaviruses have been experimentally confirmed to use DPP4 (1, 3, 4, 7, 48). The inability of most merbecoviruses to engage DPP4 has hindered the development of robust infection models, limiting our understanding of their entry mechanisms and our ability to develop and evaluate the efficacy of antiviral therapeutics and vaccines.

Recent studies demonstrated that ACE2 recognition has evolved independently in multiple merbecovirus clades with distinct geographic distributions (17, 19, 20). Our findings extend this paradigm by identifying a subset of HKU25 clade coronaviruses as group 3 ACE2-using merbecoviruses, challenging the previously proposed use of DPP4 by HKU25 clade viruses (7). These results unveil the prevalence of ACE2 usage among merbecoviruses and the overall similar but divergent ACE2 engagement mode utilized by HKU5 and HKU25 clades concurs with their close RBD phylogenetic relationships (18, 20). The recognition of partially overlapping surfaces for multiple merbecoviruses, sarbecoviruses, and setracoviruses using entirely distinct RBM architectures suggest fulfillment of specific geometric constraints leading to viral entry (49, 50).

Notably, some merbecoviruses may employ receptors other than ACE2 or DPP4. For instance, EriCoVs, NsGHA2010, and several merbecoviruses from the HKU25 clade exhibit distinct RBM indels and are not found to use any tested ACE2 or DPP4 orthologs (**Fig. S1*D***). However, while three HKU25 clade coronaviruses (VsCoV-1, VmSL2020, and VmSL2021) were not confirmed as ACE2 dependent in this study, their reliance on ACE2 cannot be excluded due to untested ACE2 orthologs from *V.mur* and *V.sin*. Similarly, polymorphisms in the *P.aur* ACE2 allele may have influenced receptor functionality for PaGB01 (51). Confirming the host ACE2/DPP4 receptor orthologs from the sampling host or identifying new receptors for these viruses will be essential to obtain a comprehensive understanding of receptor utilization and species tropism for merbecoviruses.

The evolutionary history of receptor usage among merbecoviruses remains unclear. Accumulating evidence suggests that S recombination with breaking points between NTD and S_2_ subunit plays a key role in receptor switching (14, 52). For instance, DPP4-using MERS-CoV and BtCoV-422 are proposed to have acquired their RBDs through recombination with HKU4 clade viruses from NeoCoV-like and EjCoV-3-like ACE2-using ancestors, respectively (14).

Whereas RBD recombination facilitates receptor switching in coronaviruses, this mechanism does not establish novel receptor recognition modalities. Instead, remodeling of interactions via critical adaptive sequence changes, such as RBM indels and antigenic drift, appear to drive the acquisition of new receptor binding modes (53). Although ACE2 utilization has been proposed to have emerged independently across diverse bat species, the evolutionary origins of the conserved DPP4-binding mode in merbecoviruses remain enigmatic. The limited genetic diversity and restricted geographic distribution of HKU4 clade viruses support an evolutionary trajectory involving RBM indel-driven divergence from ancestral HKU25- or HKU5-like lineages. In agreement with a prior hypothesis (14), HKU4 clade recombination with ACE2-using lineages ultimately generated phylogenetically discrete DPP4-using MERSr-CoVs including MERS-CoV and BtCoV-422 (**Fig. 6**). This evolutionary model is supported by shared distinct sequence signatures of indels 2 and 3 among DPP4-using viruses (e.g. MERS-CoV, HKU4, BtCoV-422) in **Fig. 1 *D* and *E***. The RBD phylogenetic gap between MERS-CoV and currently known HKU4 strains, alongside the absence of viruses highly similar to MERS-CoV in extensive bat virome surveys(54–60), implies the existence of unsampled reservoirs of HKU4-like viruses in uncharacterized ecological niches, which may have donated the DPP4-binding RBD sequences to MERS-CoV.

**Fig. 6.**
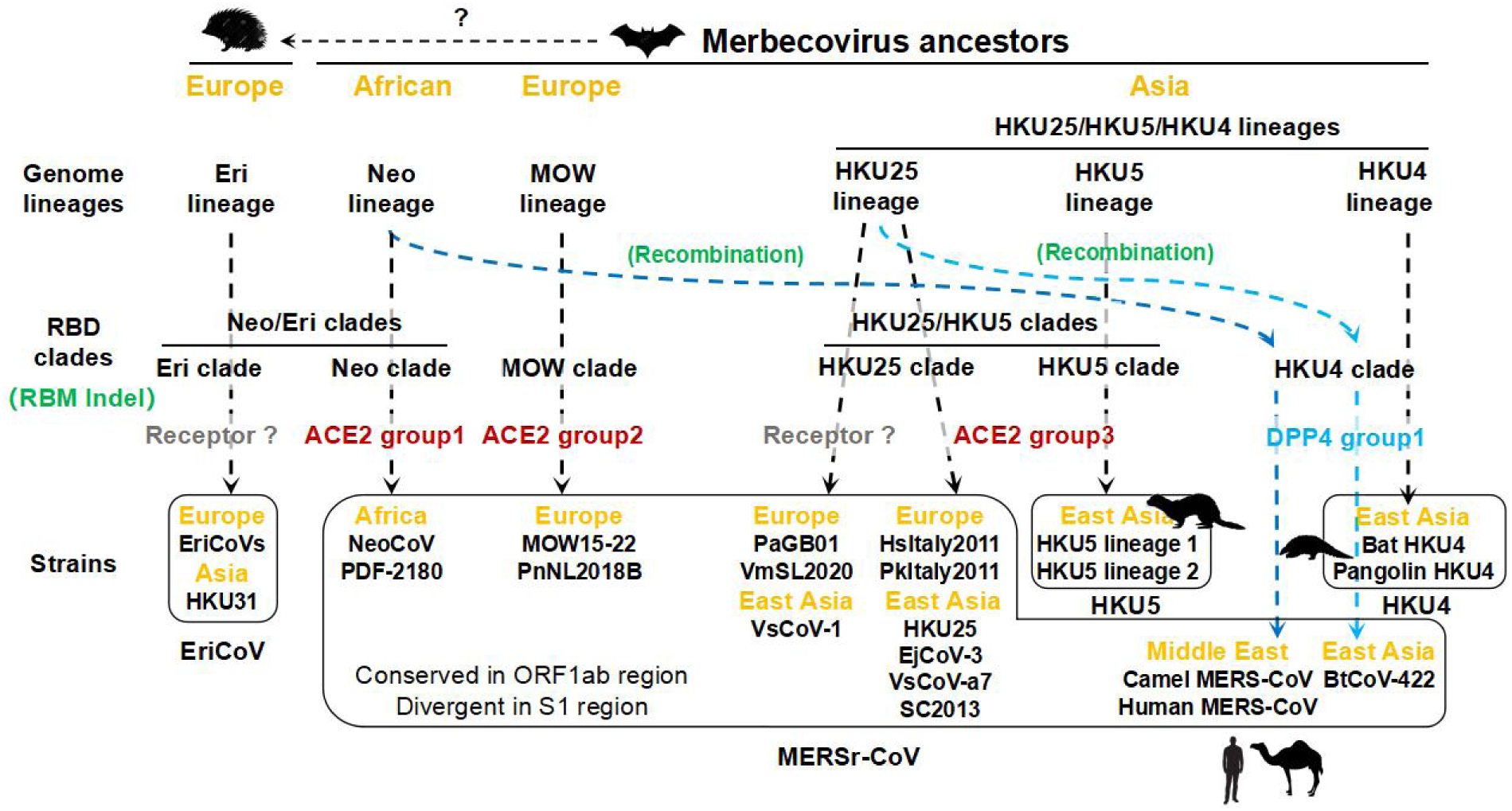
Proposed evolutionary model of merbecovirus receptor usage acquisition and switching. Geographical regions (orange) and receptor binding modes of specific merbecovirus RBD clades (gray: unidentified; red: ACE2; blue: DPP4) are indicated. Genome lineage evolution and RBD clade diversification involve RBM indels and S_1_ recombination (green) are annotated. Light and blue dashed lines propose the origins of DPP4-using MERS-CoV and BtCoV-422. Strains of the same species are grouped within black boxes. Spillovers to non-bat mammalian species are indicated. Indel: sequence insertions and deletions.

The HKU25 clade encompasses genetically diverse MERSr-CoVs circulating across vespertilionid bats in Eurasia. Host migration and cross-species transmission appear to drive viral diversification and recurrent recombination events. Despite being phylogenetically classified within MERSr-CoVs based on ORF1ab sequences, HKU25 clade coronaviruses resemble HKU5 clade viruses in terms of ACE2 binding mode and ortholog tropism. As observed for other merbecovirus clades, critical interface residues and glycans govern their ACE2 specificity and zoonotic potential (17, 19, 21). For example, the *P.abr* ACE2-specific N386-glycan, which is accommodated upon HKU5 RBD binding, appears to sterically prevent the recognition of HKU25 clade coronaviruses (**Fig. 4 *C* and *F***). Unlike HKU5, which has spilled over into minks and evolved hACE2-compatible lineages(23, 61), currently sampled HKU25 clade coronaviruses remain bat-restricted and poorly adapted to human ACE2 (hACE2). The absence of furin cleavage sites in HKU25 clade S proteins may constitute additional constraints on human adaptation. Nevertheless, the broad ACE2 ortholog tropism and hACE2 utilization of EjCoV-3 (albeit weak) reveals pre-adaptive potential for host switching of this clade and thus warrants close monitoring. Lessons learned from the COVID-19 pandemic, proactive investigations of transmissibility, pathogenicity, and therapeutic vulnerabilities of these ACE2-using merbecoviruses should be prioritized for our preparedness for potential future outbreaks.

## Materials and Methods

Cell lines, Construct design, Recombinant protein production, Cell-cell fusion assay, Immunofluorescence assay, Biolayer interferometry, Flow cytometry, rcVSV-CoV amplification and inhibition assays, Cryo-electron microscopy data collection, processing, and model building, Bioinformatic and structural analysis, and Statistical analysis are described in *SI appendix, SI Materials and Methods*. Materials generated in this study will be made available after completion of a materials transfer agreement.

### RBD-hFc live-cell binding assay

HEK293T cells transiently expressing receptors were incubated with RBD-hFc proteins (diluted in DMEM) for 30 minutes at 37°C (36 hours post-transfection). Subsequently, cells were washed once with HBSS and incubated with 1 μg/mL of Alexa Fluor 488-conjugated goat anti-human IgG (Thermo Fisher Scientific; A11013) diluted in HBSS/1% BSA for 1 hour at 37℃. After additional washing with HBSS, nuclei were stained with Hoechst 33342 (1:10,000 dilution in HBSS) for 30 minutes at 37℃. The images were captured using a fluorescence microscope (MI52-N). Relative fluorescence units (RFUs) were quantified using a Varioskan LUX Multi-well Luminometer (Thermo Scientific). Heatmap presentations in Figures 3A and 3B were plotted by subtracting background RLUs from control cells without ACE2 expression (Vector).

### Pseudovirus production and entry assays

VSV-dG pseudovirus (PSV) carrying trans-complemented S glycoproteins from various coronaviruses were produced following a modified protocol as previously described (62). Briefly, HEK293T cells were transfected with plasmids encoding S glycoproteins. At 24 hours post-transfection, cells were transduced with VSV-G trans-complemented VSV-dG encoding GFP and firefly luciferase (VSV-dG-fLuc-GFP, constructed and produced in-house) at 1.5×10^6^ TCID_50_, diluted in DMEM with 8 μg/mL polybrene, and incubated at 37 ℃ for 4-6 hours. After three washes with PBS, the culture medium was replaced with SMM 293-TII Expression Medium (Sino Biological, M293TII), along with the presence of the I1 neutralizing antibody targeting the VSV-G to eliminate background entry signal from the residual VSV-G-harboring pseudovirus. Supernatants containing S-incorporated VSV pseudovirus were harvested 24 hours later, centrifuged at 12,000 × g for 5 minutes (4°C), aliquoted, and stored at −80°C. The TCID_50_ of the pseudovirus was calculated using the Reed-Muench method (63, 64).

For single-round VSV pseudovirus entry assays, HEK293T or Caco2 cells transiently/stably expressing different receptors (3×10⁴ cells/well in 96-well plates) were incubated with pseudovirus (2×10^5^ TCID_50_/100 μL). Pseudoviruses produced in serum-free SMM 293-TII Expression Medium were typically pretreated with TPCK-trypsin (Sigma-Aldrich, T8802) for 10 minutes at room temperature, followed by 10% FBS in the culture medium to inactive the protease activity. The I1-neutralizing antibody was added to the trypsin-treated pseudoviruses again to reduce the background before use. Luciferase activity (Relative light units, RLU) was measured at 18 hpi using the Bright-Glo Luciferase Assay Kit (Promega, E2620) and detected with a GloMax 20/20 Luminometer (Promega) or Varioskan LUX Multi-well Luminometer (Thermo Fisher Scientific).

To examine the S glycoprotein packaging and cleavage efficiency, the S-incorporated VSV pseudoviruses were concentrated using a 30% sucrose cushion (30% sucrose, 15 mM Tris-HCl, 100 mM NaCl, 0.5 mM EDTA) at 20,000 × g for 1 hour at 4℃. Pellets were resuspended in 1×SDS loading buffer, vortexed, boiled (95°C, 10 minutes), and followed by western blot detecting the S glycoproteins by C-terminal HA tags and with the VSV-M serving as a loading control.

## Data availability

The cryo-EM maps and model have been deposited to the electron microscopy data bank and protein data bank with accession numbers EMD-49092, PDB-9N7D (*R.nor* ACE2-bound HsItaly2011 RBD), and EMD-49093, PDB-9N7E (*E.fus* ACE2-bound VsCoV-a7 RBD). Any additional information required to reanalyze the data reported in this paper is available from the lead contact upon request.

## Acknowledgments

This study was supported by National Natural Science Foundation of China (NSFC) projects (82322041, 32270164 to H.Y., 323B2006 to C.-B.M.), the National Key R&D Program of China (2023YFC2605500 and 2023YFC2607300 to H.Y.), Natural Science Foundation of Hubei Province (2023AFA015 to H.Y.), the Fundamental Research Funds for the Central Universities (to H.Y.) and TaiKang Center for Life and Medical Sciences (to H.Y.). Yan Lab thanks Lu Lu (Fudan University) for providing EK1C4 peptides; Qiang Ding (Tsinghua University), Qihui Wang (CAS Key Laboratory of Pathogenic Microbiology & Immunology, China), and Zheng-Li Shi (Guangzhou National Laboratory) for sharing some ACE2 plasmids with H.Y. used in this study.

This study was also supported by the National Institute of Allergy and Infectious Diseases (P01AI167966, DP1AI158186 and 75N93022C00036 to D.V.), a Shurl and Kay Curci Foundation Graduate Scholarship Award (to R.G.), the National Institute of General Medical Sciences, an Investigators in the Pathogenesis of Infectious Disease Awards from the Burroughs Wellcome Fund (D.V.), the University of Washington Arnold and Mabel Beckman cryo-EM center and the National Institute of Health grant S10OD032290 (to D.V.). D.V. is an Investigator of the Howard Hughes Medical Institute and the Hans Neurath Endowed Chair in Biochemistry at the University of Washington.

## Author contributions

C.L., Y.-J.P., D.V., and H.Y. conceived the project. Y.-J.P. and C.S. designed glycoprotein constructs and recombinantly expressed glycoproteins. C.L., C.-B.M., Y.-C.S., and M.Y.L. cloned S, RBD-hFc, and ACE2 mutants and conducted RBD-hFc binding assays. C.L. and C.-B.M. conducted S cleavage and cell-cell fusion assays. C.L. and X.Y. rescued the rcVSV-HKU25r-S pseudotypes and C.L. performed rcVSV propagation and inhibition assays. C.-B.M., C.S. and R.G. conducted biolayer interferometry binding experiments. C.L., C.-B.M., and Y.-C.S. carried out VSV pseudovirus entry and neutralization assays. Y.-J.P. carried out cryo-EM sample preparation, data collection, and processing. Y.-J.P. and D.V. built and refined the structures. DV and HY wrote the manuscript with input from all authors. H.Y., D.V., C.L., Y.-J.P., C.S., R.G., and C.-B.M. analyzed the data. C.L. conducted phylogenetic and conservation analysis.

## Declaration of interests

The authors declare no competing interests.

## SUPPLEMENTARY INFORMATION

### Materials and Methods

#### Cell lines

HEK293T (CRL-3216), HEK293T (ATCC, CRL-11268), Caco2 (HTB-37), BHK-21 (CCL-10), and I1-Hybridoma (CRL-2700) cells were obtained from the American Type Culture Collection (ATCC). Expi293F (Thermo Fisher Scientific, A14527) was used for protein production. All the above cells were cultured in Dulbecco’s Modified Eagle Medium (DMEM, Monad, China) supplemented with 1% PS (Penicillin/Streptomycin) and 10% Fetal Bovine Serum (FBS). The I1-Hybridoma cell line, which produces a neutralizing antibody targeting the VSV glycoprotein (VSV-G), was maintained in Minimum Essential Medium (MEM) with Earles’s balances salts, 2.0 mM of L-glutamine (Gibico), and 10% FBS. All cell lines were incubated at 37℃ with 5% CO2 and routinely passaged every 2-3 days. HEK293T or Caco-2 cell lines overexpressing various receptors were generated using lentivirus transduction followed by the puromycin (1 μg /ml) selection.

#### Construct design

Plasmids of WT or mutated mammalian ACE2 orthologs or ACE2 chimera were constructed by inserting human codon-optimized coding sequences into the lentiviral transfer vector (pLVX-EF1a-Puro, Genewiz) with C-terminus 3 × FLAG tags (DYKDHD-G-DYKDHD-I-DYKDDDDK) for bat ACE2 and single FLAG tags (DYKDDDDK) for non-bat mammalian ACE2 orthologs (21, 65, 66). The constructions expressing human or bat DPP4 orthologs encoding residues 1 to 766 corresponding to hDPP4 were generated similarly to ACE2 orthologs with 3 × FLAG tags. For S-incorporated VSV pseudovirus production, human codon-optimized S sequences from MERS-CoV (YP_009047204.1), HKU5-1 (YP_001039962), HKU5-19S (AGP04932.1), HKU25-NL140462 (ASL68953.1), HKU25-NL13892 (AWH65943.1), EjCoV-3 (BDD37132.1), SC2013 (AHY61337.1), VsCoV-kj15 (BDI08820.1), VsCoV-a7 (BDI08829.1), HsItaly2011 (AUM60014.1), VmSL2020 (USF97409.1), VsCoV-1 (BBJ36008.1), PaGB01 (WDQ26963.1) were cloned into the pCAGGS vector with C-terminal residues 13–15 replaced by a HA tag (YPYDVPDYA) to facilitate S incorporation (67). For the expression of recombinant CoVs RBD-hFc fusion proteins, plasmids were constructed by inserting RBD coding sequences from HKU5-19s (residues 385-586), HKU25-462 (residues 385-587), HKU25-892 (residues 384-586), HKU25-305 (residues 385-587), PaGD2016-Q249 (residues 383-584), EjCoV-3 (residues 383-585), SC2013 (residues 382-588), VsCoV-kj15 (residues 388-594), HsItaly2011 (residues 383-586), PkItaly2011 (residues 383-586), VsCoV-a7 (residues 385-587), VmSL2020 (residues 389-590), VmSL2021 (residues 387-588), VsCoV-1 (residues 382-583), and PaGB01 (residues 377-572) into the pCAGGS vector containing an N-terminal CD5 secretion signal peptide (MPMGSLQPLATLYLLGMLVASVL) and C-terminal hFc-twin-strep-3 × FLAG tags (WSHPQFEKGGGSGGGSGGSAWSHPQFEK-GGGRSDYKDHDGDYKDHDIDYKDDDDK) for purification and detection. Plasmids expressing soluble ACE2 ectodomain proteins were generated by inserting sequences from hACE2 (residues 18-740), *E.fus* ACE2 (residues 18-746), and *R.nor* ACE2 (residues 18-740) into the pCAGGS vector, with an N-terminal CD5 secretion signal peptide and a C-terminal twin-strep-3 × FLAG tag. DNA fragments for cloning chimera or mutants were generated by overlap extension PCR or gene synthesis and verified by commercial DNA sequencing.

For cryo-EM analysis, the ACE2 ectodomain encoding residues of *R.nor* (1–741) and *E.fus* (1–747) were subcloned into the pcDNA3.1(+) plasmids with C-terminal Avi and octa-histidine tag. The RBD encoding residues of HsItaly2011 (388–589) and VsCoV-a7 (356–556) were subcloned into the pcDNA3.1(+) with N-terminal signal peptide (MGILPSPGMPALLSLVSLLSVLLMGCVAETGT) and C-terminal thrombin cleavage sequence, 8 flexible GS linker sequence and an Avi tag followed by octa-histidine tag.

#### Recombinant protein production

For producing proteins or antibodies for biochemical or neutralization assays, corresponding plasmids expressing proteins were transfected using GeneTwin reagent (Biomed, TG101-01) in HEK293T cells or Expi293F cells. After 4-6 hours post-transfection, culture medium was replenished with the SMM 293-TII Expression Medium (Sino Biological, M293TII). Protein-containing supernatant was collected every three days for 2-3 batches. Fc-tagged proteins (Antibodies and recombinant RBD-hFc) were purified using Pierce Protein A/G Plus Agarose (Thermo Scientific, 20424). In general, proteins were enriched by the agarose, washed with wash buffer (100 mM Tris/HCl, pH 8.0, 150 mM NaCl, 1 mM EDTA), eluted using the Glycine buffer (100 mM in H_2_O, pH 3.0), and immediately neutralized with 1/10 volume of 1M Tris-HCI, pH 8.0 (15568025, Thermo Scientific). Proteins with twin-strep tag were purified using Strep-Tactin XT 4Flow high-capacity resin (IBA, 2-5030-002), washed with wash buffer (100 mM Tris/HCl, pH 8.0, 150 mM NaCl, 1 mM EDTA), and then eluted with buffer BXT (100 mM Tris/HCl, pH 8.0, 150 mM NaCl, 1 mM EDTA, 50 mM biotin). Purified proteins were concentrated using ultrafiltration tubes, buffer-changed to PBS, and stored at -80℃. Concentrations were determined by the Omni-Easy Instant BCA Protein Assay Kit (Epizyme, ZJ102).

For recombinant glycoprotein production for cryo-EM analysis, each construct was expressed in Expi293F cells (Thermo Fischer Scientific), cultured at 37°C with constant rotation at 130 RPM in a humidified incubator with 80% relative humidity and 8% CO2. DNA was transfected following the protocol outlined by the manufacturer (Thermo Fischer Scientific) and grown for four days prior to harvest. Cell culture supernatants were clarified by centrifugation and harvested using either HisTrap HP Ni Sepharose Columns (Cytiva) or Ni Sepharose excel resin (Cytiva). The resin was washed with 10-50 CVs of 25 mM Tris 150 mM NaCl 10 mM Imidazole pH 8.0, followed by a 15 CVs wash using 25 mM Tris 150 mM NaCl 400 mM Imidazole pH 8.0 to elute the protein. Afterwards, the proteins were buffer exchanged into 25 mM Tris 150 mM NaCl pH 8.0 using 10KDa or 100KDa Amicom Ultra-15 Centrifugal Filter Units (Millipore) for RBDs or ACE2s respectively. A portion of proteins were also set aside and biotinylated using a biotin ligase (BirA) reaction kit (Avidity). These biotinylated RBDs were adjusted to a final concentration of 40 μM with all the provided reagents and the reaction was carried out at room temperature for 30 minutes followed by 10 hours at 4°C. Subsequently, proteins were each purified by gel filtration using a Superose-6 Increase 10/300 column (ACE2s) or Superdex-200 10/300 column (RBDs) (Cytiva) equilibrated in a buffer containing 25 mM Tris, 150 mM NaCl pH 8.0. The main peak was collected, flash-frozen using liquid nitrogen, and stored at -80°C until use.

#### Cell-cell fusion assays

Dual-split proteins (DSPs) based fusion assays were performed in Caco-2 cells stably expressing ACE2 receptors. Two group cells stably expressing the indicated receptors were transiently transfected with different plasmids for assessing the HKU25-related coronaviruses S and receptor interaction-mediated membrane fusion. Group A cells were transfected with plasmids encoding S and rLucN(1–155)-sfGFP1-7(1–157), while group B cells were transfected with plasmids encoding S and sfGFP8-11 (158–231)-rLuc (156–311) expressing plasmids. At 12 hours post-transfection, two groups of cells were trypsinized, mixed, and seeded into a 96-well plate at 8 × 10^4^ cells per well. At 24 hours post-transfection, the cells were washed once with DMEM and then incubated with DMEM with or without indicated concentrations of TPCK-treated trypsin (Sigma-Aldrich, T8802) for 10 minutes at room temperature. After washing with DMEM, the cells were replenished with DMEM/10% FBS to neutralize trypsin activity. Syncytia formation with green fluorescence was assessed 6 hours later using Hoechst 33342 nuclear staining (1:5,000 dilution in Hanks’ Balanced Salt Solution (HBSS) for 30 minutes at 37 °C) and fluorescence microscopy (MI52-N; Mshot).

#### Immunofluorescence assay

For assessing the expression levels of ACE2 or DPP4 orthologs tags, cells transiently or stably expressing the indicated receptors with C-terminal fused FLAG tags were fixed and permeabilized by incubation with 100% methanol (10 minutes at room temperature), washed by HBSS, and incubated with a mouse antibody M2 (Sigma-Aldrich, F1804) diluted in PBS/1% BSA for one hour at 37℃. After one HBSS wash, the cells were incubated with Alexa Fluor 594-conjugated goat anti-mouse IgG (Thermo Fisher Scientific, A32742) secondary antibody diluted in 1% BSA/PBS for one hour at 37℃. The images were captured with a fluorescence microscope (Mshot, MI52-N) after the nuclei were stained with Hoechst 33342 reagent (1:1,000 dilution in HBSS).

#### Biolayer interferometry (BLI)

For dimeric hACE2 ectodomain proteins binding to immobilized HKU25-NL13892 RBD-hFc or HsItaly2011 RBD-hFc, recombinant RBD-hFc proteins were immobilized on Protein A (ProA) biosensors (ForteBio, 18-5010), which were then incubated with the indicated soluble Dimeric hACE2-ectodomain proteins (two-fold serial-diluted in PBST starting from 2,000 nM or 1000 nM) with wells incubated with kinetic buffer (PBST) only as a background control. Protein binding kinetics was assessed using an Octet RED96 instrument (Molecular Devices) at 25℃ and shaking at 1,000 rpm. The kinetic parameters and the apparent binding affinities (due to ACE2 dimerization) were analyzed using Octet Data Analysis software 12.2.0.20 with global curve fitting using a 1:1 binding model.

Biotinylated HsItaly2011 and VsCoV-a7 RBD were diluted into 10x Octet Kinetics Buffer (Sartorius) and loaded onto pre-hydrated streptavidin biosensors to a 1 nm shift. The tips were then re-equilibrated in the kinetics buffer before being dipped into a serial dilution of *R.nor* or *E.Fus* ACE2 dimers for 300 to 500 seconds followed by another incubation in kinetics buffer to assess the dissociation. The ACE2 starting concentrations were as high as 3,000 nM to as low as 900 nM, and diluted either two or three-fold in the kinetics buffer leaving one well without any dilution as a background control. Kinetics were assessed at 30℃ and 1,000 rpm using an Octet Red96. The binding kinetics were baseline subtracted and assessed using Octet Data Analysis 11.1 software with a global curve fitting in a 1:1 binding model and plotted in GraphPad 10.4.

#### Flow cytometry

Cells transiently expressing ACE2 or DPP4 orthologs were washed twice with cold PBS and incubated with 10 μg/mL indicated RBD hFc proteins at 4℃ for 30 minutes at 36 hours post-transfection. Subsequently, cells were incubated with Alexa Fluor 488-conjugated goat anti-human IgG to stain the bound RBD-hFc (Thermo Fisher Scientific; A11013) at 4℃ for 1 hour. Subsequently, cells were detached with 5 mM EDTA/PBS, fixed with 4% PFA, permeabilized with 0.25% Triton X-100, blocked with 1% BSA/PBS at 4℃, and then incubated with mouse anti-FLAG tag antibody M2 (Sigma-Aldrich, F1804) diluted in PBS/1% BSA for 1 hour at 4℃, followed by incubation with Alexa Fluor 647-conjugated goat anti-mouse IgG (Thermo Fisher Scientific; A32728) diluted in 1% BSA/PBS for 1 hour at 4℃. For all samples, 10,000 receptor-expressing live cells (gated based on SSC/FSC and FLAG-fluorescence intensity and SSC/FSC) were analyzed using a CytoFLEX Flow Cytometer (Beckman).

#### rcVSV-CoV amplification and inhibition assays

The experiments of replication-competent VSV-S (rcVSV-S) were authorized by the Biosafety Committee of the State Key Laboratory of Virology and Biosafety, Wuhan University, and conducted under BSL2 conditions. To construct plasmids for rescuing replication-competent (rc) VSV-CoV expressing HKU25-clade S glycoproteins, the firefly luciferase (fLuc) encoding sequences of pVSV-dG-fLuc-GFP (50) were replaced with the indicated coronavirus spike sequences. Reverse genetics was applied to rescue rcVSV-CoV-S pseudotypes expressing HKU25-clade S glycoproteins along with a GFP reporter, following a modified protocol from previous descriptions (62). Briefly, BHK-21 cells were seeded in a 6-well plate at 80% confluence and inoculated with 5 MOI of recombinant vaccinia virus expressing T7 RNA polymerase (vvT7, a kind gift from Mingzhou Chen’s lab, Hubei University) for 45 minutes at 37°C. Subsequently, cells were transfected with pVSV-dG-GFP-S vector plasmids and helper plasmids (pVSV-dG-GFP-S: pBS-N: pBS-P: pBS-G: pBS-L=5:3:5:8:1) after washing by DMEM. The rcVSV-CoV containing supernatant (P0) was filtered (0.22 μm) and amplified in Caco-2 cells transiently expressing VSV-G (P1). Subsequently, P2 viruses were generated in Caco-2 cells stably expressing indicated ACE2, without the ectopic expression of VSV-G and in the presence of anti-VSVG antibody (I1-Hybridoma supernatant) to produce viruses without VSV-G contamination. For amplification assay, 3×10^4^ trypsinized Caco-2 cells stably expressing the indicated ACE2 were incubated with rcVSV-CoV (1×10^4^ TCID_50_/100 μL) in a 96-well plate in DMEM supplemented with 2% FBS with or without the treatment of indicated concentrations of TPCK-treated trypsin. At the indicated time post-infection, the cell nuclei were stained with Hoechst 33342 (1:10,000 dilution in HBSS) for 30 minutes at 37℃, and the fluorescence images were taken by a fluorescence microscope (MI52-N).

#### Cryo-electron microscopy data collection, processing, and model building

The *E.fus* ACE2 ectodomain-bound VsCoV-a7 RBD complex was prepared by mixing at 1:1.2 molar ratio followed by a 1 hour incubation at room temperature. 3µL of 5 mg/ml complex with 6 mM 3-[(3-Cholamidopropyl)dimethylammonio]-2-hydroxy-1-propanesulfonate (CHAPSO) were applied onto freshly glow discharged R 2/2 UltrAuFoil grids(68) prior to plunge freezing using a vitrobot MarkIV (ThermoFisher Scientific) with a blot force of 0 and 5.5 sec blot time at 100% humidity and 22°C. The data was acquired using an FEI Titan Krios transmission electron microscope operated at 300 kV and equipped with a Gatan K3 direct detector and Gatan Quantum GIF energy filter, operated in zero-loss mode with a slit width of 20 eV. Automated data collection was carried out using Leginon(69) at a nominal magnification of 105,000× with a pixel size of 0.843 Å. The dose rate was adjusted to 9 counts/pixel/s, and each movie was acquired in counting mode fractionated in 100 frames of 40 ms. A total 14,595 micrographs were collected with a defocus range between -0.2 and -3 μm. Movie frame alignment, estimation of the microscope contrast-transfer function parameters, particle picking, and extraction were carried out using Warp(70). Particles were extracted with a box size of 192 pixels with a pixel size of 1.686Å. Two rounds of reference-free 2D classification were performed using cryoSPARC(71) to select well-defined particle images. Initial model generation was carried out using ab-initio reconstruction in cryoSPARC and the resulting maps were used as references for heterogeneous 3D refinement. Particles belonging to classes with the best resolved RBD and ACE2 density were selected. To further improve the data, the Topaz model(72) was trained on Warp-picked particle sets belonging to the best classes after 2D classification and particles picked using Topaz were extracted and subjected to 2D-classification and heterogenous 3D refinements. The two different particle sets from the Warp and Topaz picking strategies were merged and duplicates were removed using a minimum distance cutoff of 90Å. After two rounds of ab-initio reconstruction-heterogeneous refinements, 3D refinement was carried out using non-uniform refinement in cryoSPARC(73). The dataset was transferred from cryoSPARC to Relion using the pyem program package and particle images were subjected to the Bayesian polishing procedure implemented in Relion(74) during which particles were re-extracted with a box size of 320 pixels and a pixel size of 1.0 Å. To further improve the map quality, ab-initio reconstruction in cryoSPARC was used to classify the data in three bins and the generated models were used as references for heterogeneous 3D refinement. The final 3D refinements of the RBD bound ACE2 peptidase dimer structure were carried out using non-uniform refinement along with per-particle defocus refinement in cryoSPARC to yield the reconstruction at 2.7 Å resolution comprising 941,663 particles. To further improve the density of the RBD and ACE2 domain interface, the particles were symmetry expanded and subjected to focus 3D classification without refining angles and shifts using a soft mask encompassing the RBD and ACE2 domain interface using a tau value of 40 in Relion. Particles belonging to classes with the best resolved RBD-ACE2 domain interface density were selected and then subjected to local refinement using CryoSPARC. The final dataset contained 578,871 asymmetric units used for the final local refinement with a soft mask comprising one ACE2 peptidase domain and the bound RBD resulting in a 2.5 Å resolution reconstruction. Reported resolutions are based on the gold-standard Fourier shell correlation (FSC) of 0.143 criterion and Fourier shell correlation curves were corrected for the effects of soft masking by high-resolution noise substitution(75, 76). Local resolution estimation, filtering, and sharpening were carried out using cryoSPARC.

For the *R.nor* ACE2 ectodomain bound HsItaly2011 RBD structure, The complex was prepared by mixing at 1:1.2 molar ratio followed by a 1 hour incubation at room temperature. 3 µL of 5.3 mg/ml complex with 6 mM CHAPSO were applied onto freshly glow discharged R 2/2 UltrAuFoil grids prior to plunge freezing using a vitrobot MarkIV (ThermoFisher Scientific) with a blot force of 0 and 5.5 sec blot time at 100% humidity and 22°C. 8,298 movies were collected with a defocus range comprised between -0.2 and -3.0 μm. The overall data processing methods were the same as that for the E.fus ACE2 ectodomain-bound VsCoV-a7 RBD complex. The final dataset contained 831,432 asymmetric units used for the final local refinement with a soft mask comprising one R.nor ACE2 peptidase domain and the bound HsItaly2011 RBD resulting in a 2.5 Å resolution reconstruction. More details are shown in **Figures S5 and S6**.

UCSF Chimera(77), Coot(78), AlphaFold3(79), and Phenix(80) were used to fit, build, and refine the model using the sharpened and unsharpened cryo-EM maps. Validation used Phenix(80), Molprobity(81), EMRinger(82) and Privateer(83).

#### Bioinformatic and structural analysis

Merbecovirus S glycoprotein sequences were retrieved from the NCBI Virus database (https://www.ncbi.nlm.nih.gov/labs/virus/vssi/#/) on 9th August 2024. A total of 3,308 unique Betacoronavirus S glycoprotein entries was obtained through search terms “Betacoronavirus” or “BetaCoV” with advanced filters “sequence length between 1,200-1,400” and “exclude SARS-CoV-2”. Phylogenetic analysis in Geneious software identified merbecovirus sequences, which were refined to 150 non-redundant entries after excluding over-sampled MERS-CoV strains (retaining two representatives: one human-derived, one camel-derived). Subsequent NCBI BLAST searches using MERSr-CoV S protein sequences identified two additional MERS-related coronaviruses (PDF-2180 and VsCoV-1), yielding a final dataset of 152 merbecoviruses (sources and accession numbers in **Supplementary Data 1**). The RBM sequences in Fig. 1D and Fig. S1D were aligned via MAFFT with manual adjustments to optimize indel positioning. Fully and partially conserved residues were highlighted with red and green backgrounds, respectively. The RBD sequences used for evolutionary analysis in Fig. 1A were aligned via MAFFT-DASH. Phylogenetic trees were generated with IQ-Tree (version 2.0.6) using a Maximum Likelihood model with 1000 bootstrap replicates. Pairwise sequence identities were calculated in Geneious Prime (https://www.geneious.com/) following MAFFT alignment. The structures of HKU25-NL13892, SC2013, HsItaly2011, PaGB01 and HKU31 RBDs were predicted using AlphaFold3(79). Experimentally resolved structures included the NeoCoV RBD– Pipistrellus pipistrellus ACE2 complex (PDB: 7WPO), MERS-CoV RBD (4KQZ), HKU4 RBD (4QZV), HKU5 RBD (9D32), and MOW15-22 RBD (9C6O). All these RBDs were visualized and analyzed in ChimeraX (v1.7.1).

#### Statistical analysis

Most experiments were performed 2–3 times with 3 biological repeats unless otherwise specified. Representative results were shown as means ± SD as indicated in the figure legends. Unpaired two-tailed t-tests were conducted for statistical analyses using GraphPad Prism 10. *P* < 0.05 was considered significant. \**P* < 0.05,\*\**P* < 0.01, \*\*\**P* < 0.005, and \*\*\*\**P* < 0.001.

---

**Fig. S1.**
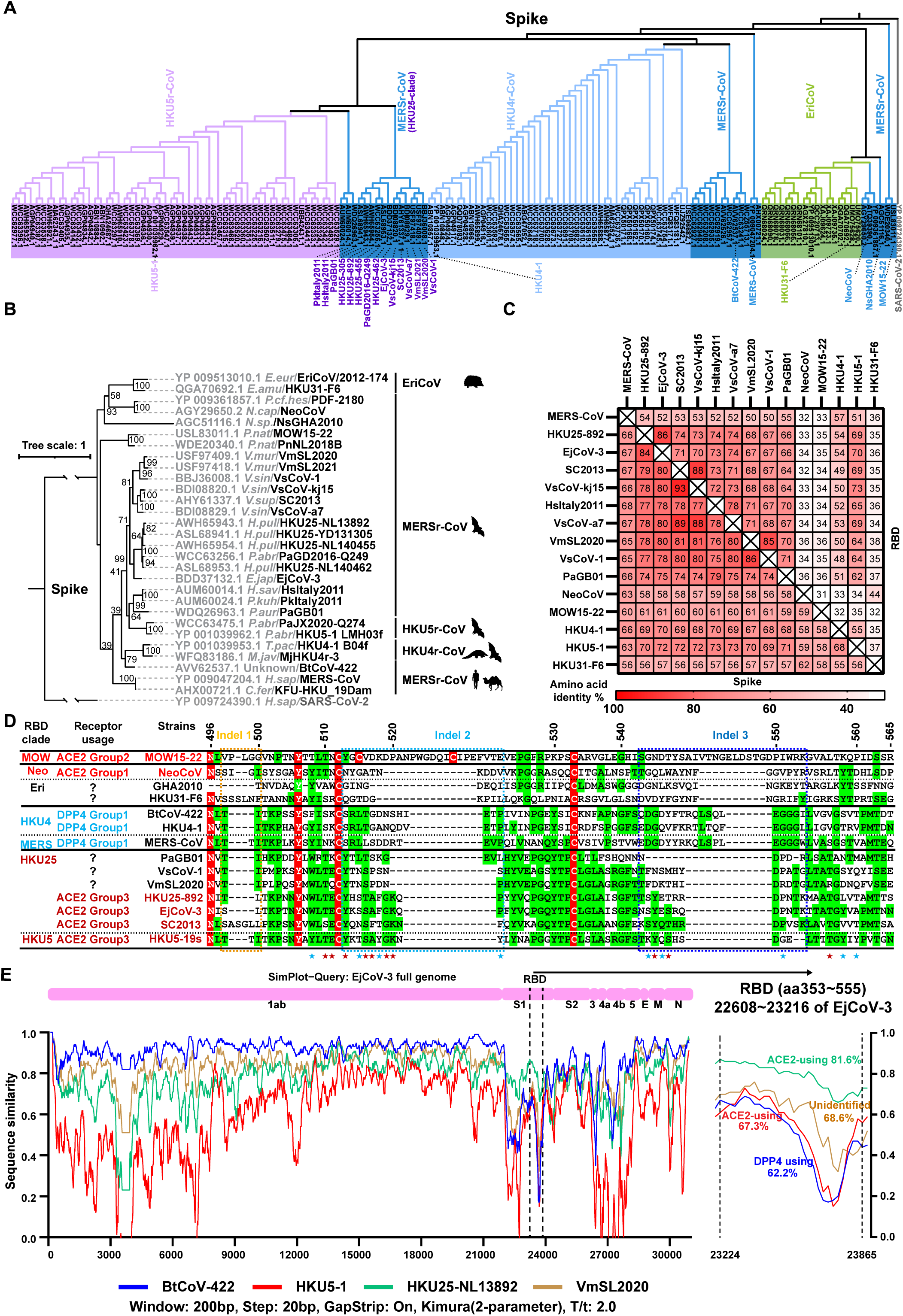
Amino acid sequence analysis of merbecovirus S glycoproteins. (***A*-*B***) phylogenetic trees based on amino acid sequences of S glycoproteins from all retrieved non-redundant merbecoviruses (A) or selected representative merbecoviruses S (B) were generated by IQ-tree2. SARS-CoV-2 was set as an outgroup. (**C**) Heat plot of pairwise RBD and S amino acid sequence identities of indicated merbecoviruses. (**D**) Manually adjusted multiple sequence alignment of RBM residues 496-565 (HKU5-19s residue numbering) from the indicated merbecoviruses with the three indels marked by dashed boxes. Fully and partially conserved residues were highlighted with red and green background, respectively. (**E**) SimPlot analysis of whole-genome nucleotide similarity of indicated merbecoviruses relative to EjCoV-3. The right panel magnifies the RBD region (EjCoV-3 positions 22608∼23216 nt). Dashed lines indicate RBD boundaries; nucleotide identities to EjCoV-3 are labeled.

**Fig. S2.**
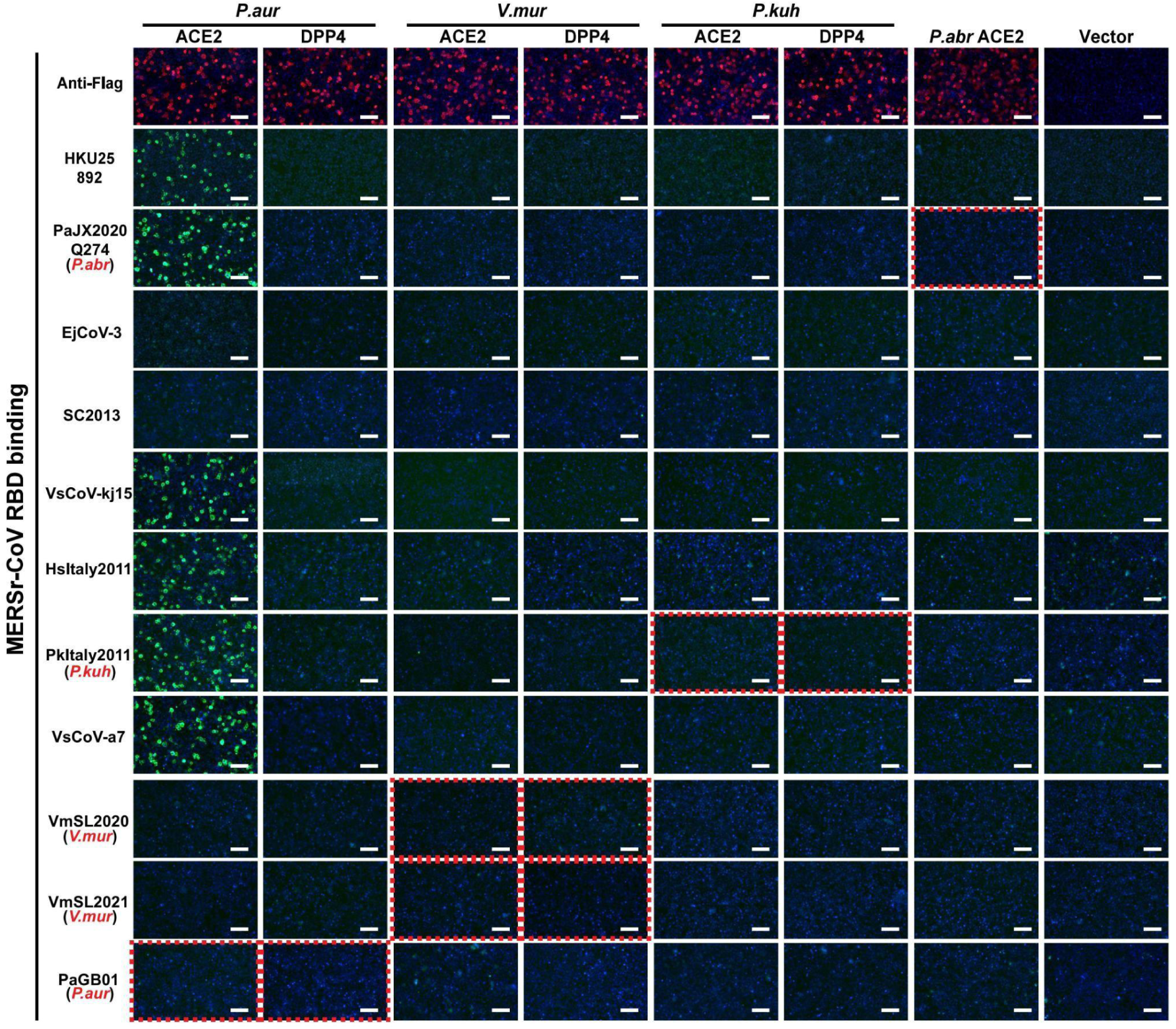
Binding of HKU25 clade RBD-hFc constructs to ACE2 or DPP4 orthologs from selected bat host species. HKU25-clade coronaviruses RBD-hFc binding to HEK293T cells transiently expressing the indicated receptors assessed by immunofluorescence. The Red dashed boxes highlight data that shows the RBD binding of indicated viruses with receptors from their reported host species (marked in red). Scale bars: 100 μm.

**Fig. S3.**
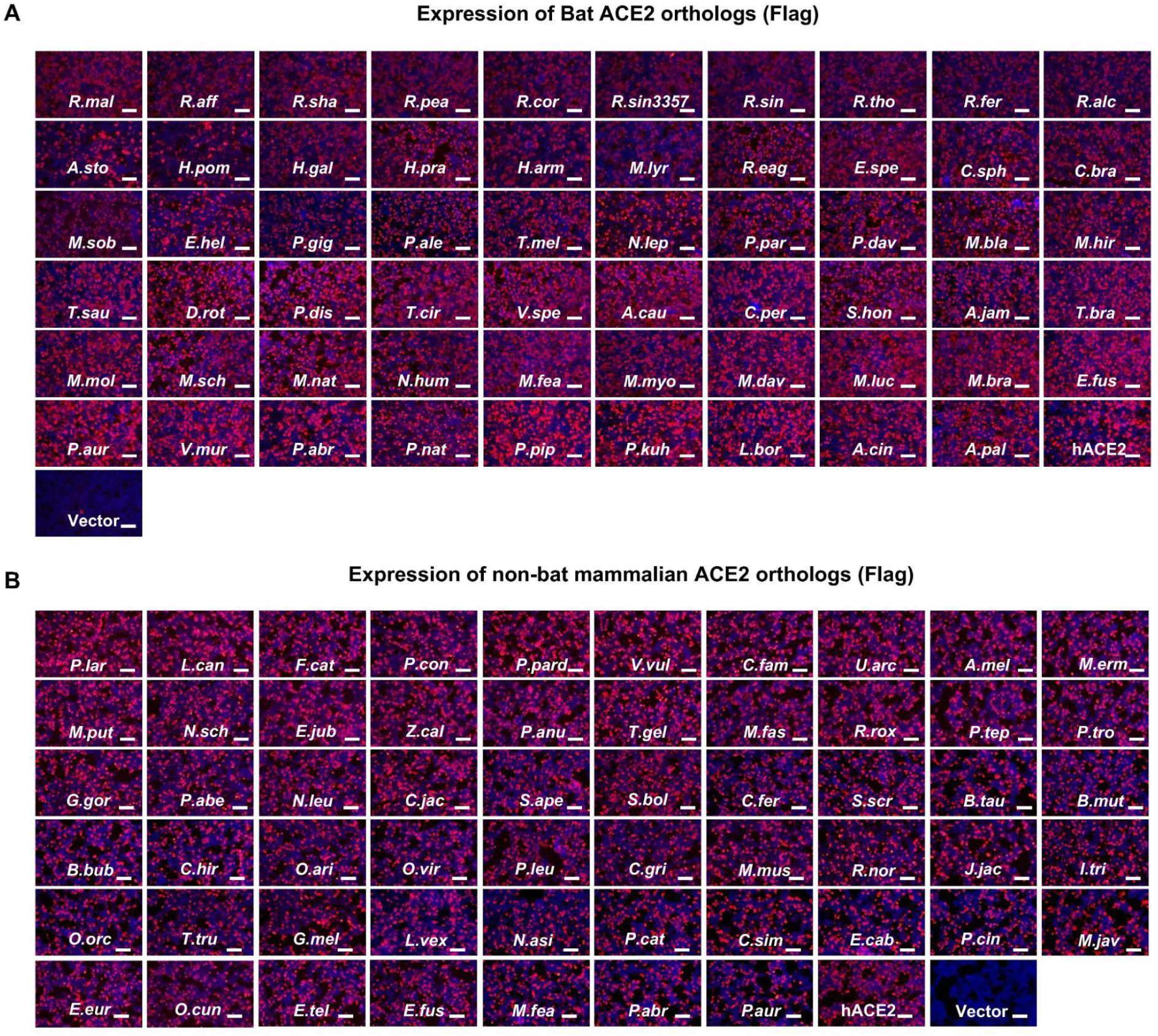
The expression level of ACE2 orthologs from various mammalian species. (***A*-*B***) Immunofluorescence analysis of the expression of bat (*A*) or non-bat mammalian (*B*) ACE2 orthologs in HEK293T cells by detecting the C-terminal fused FLAG tags.Scale bars: 100 μm.

**Fig. S4.**
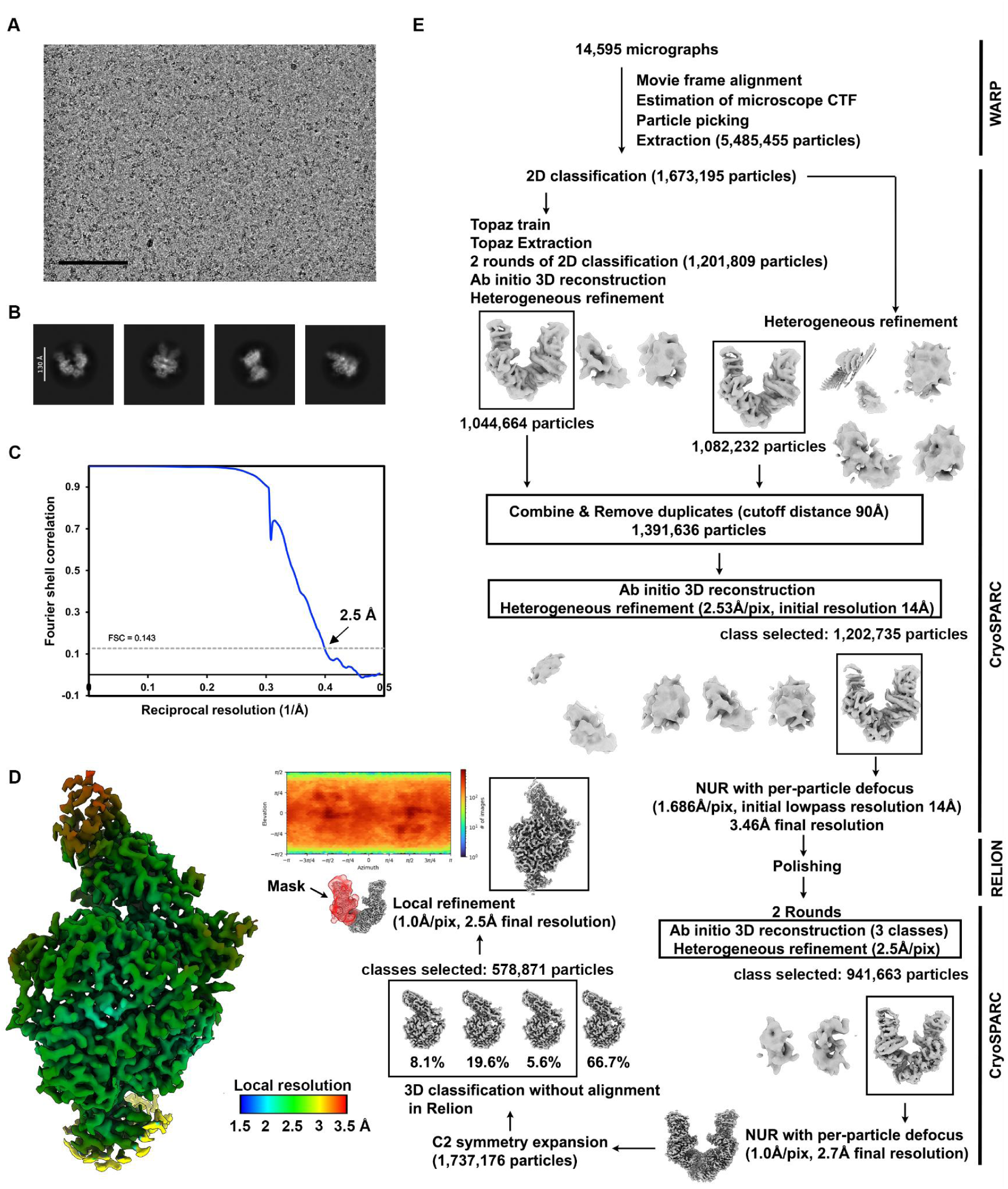
Cryo-EM data processing of the *E.fus* ACE2 bound VsCoV-a7 RBD data set. (***A*-*B***) Representative electron micrographs (A) and 2D class averages (*B*) of the *E.fus* ACE2-bound VsCoV-a7 RBD complex embedded in vitreous ice. Scale bars: 100 nm (*A*) and 130 Å (B). (***C***) Gold-standard Fourier shell correlation curve of the E.fus ACE2-bound VsCoV-a7 RBD reconstruction. The 0.143 cutoff is indicated by a horizontal dashed line. (**D**) Local resolution estimation calculated using cryoSPARC and plotted on the sharpened map. (**E**) Data processing flowchart. CTF: contrast transfer function; NUR: non-uniform refinement. The angular distribution of particle images calculated using cryoSPARC is shown as a heat map.

**Fig. S5.**
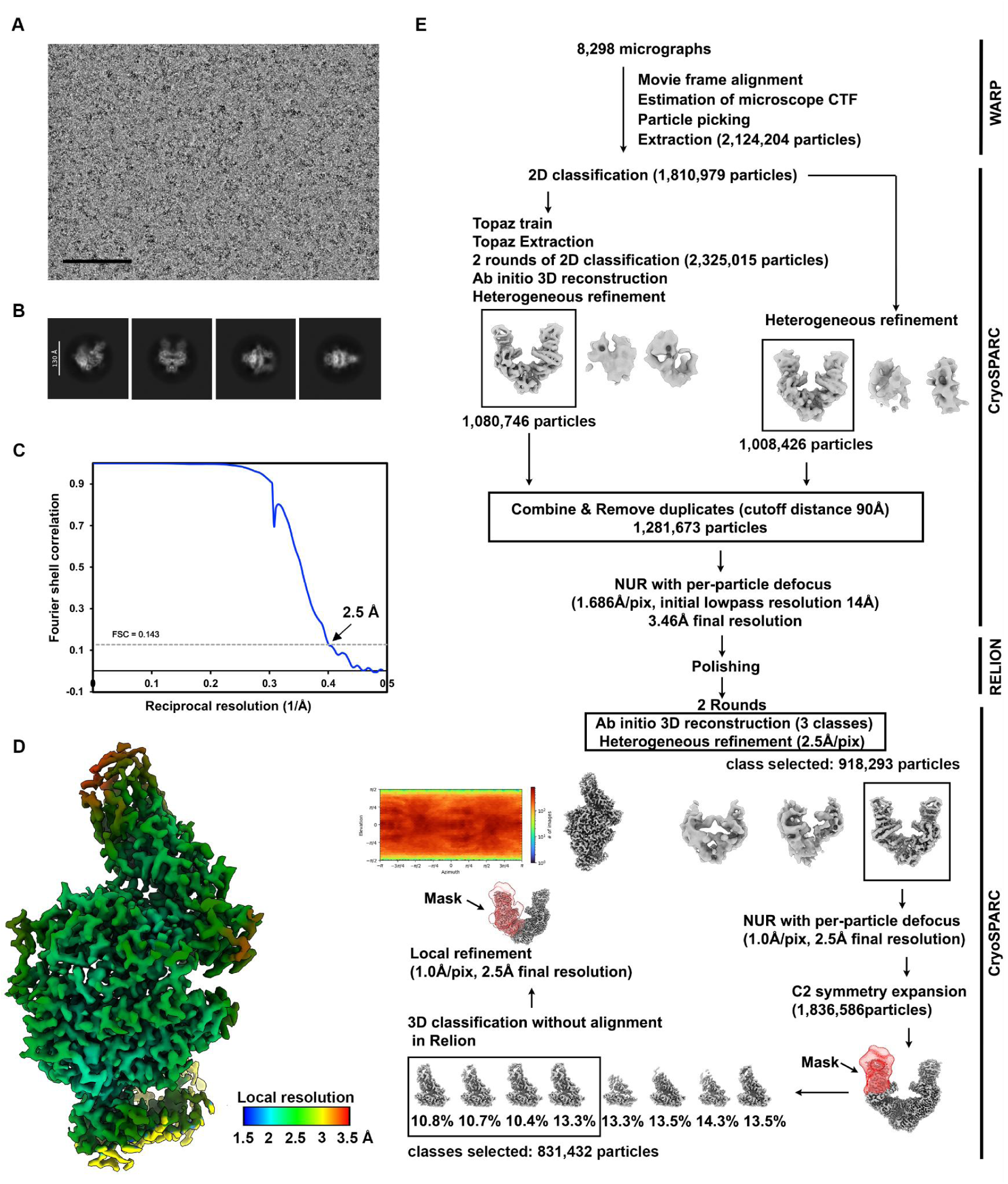
Cryo-EM data processing of the *R.nor* ACE2 bound HsItaly2011 RBD data set. (***A*-*B***) Representative electron micrographs (*A*) and 2D class averages (*B*) of the R.nor ACE2-bound HsItaly2011 RBD complex embedded in vitreous ice. Scale bars: 100 nm (A) and 130 Å (B). (***C***) Gold-standard Fourier shell correlation curve of the *R.nor* ACE2-bound HsItaly2011 RBD reconstruction. The 0.143 cutoff is indicated by a horizontal dashed line. (***D***) Local resolution estimation of the R.nor ACE2-bound HsItaly2011 RBD reconstruction calculated using cryoSPARC and plotted on the sharpened map. (***E***) Data processing flowchart. CTF: contrast transfer function; NUR: non-uniform refinement. The angular distribution of particle images calculated using cryoSPARC is shown as a heat map.

**Fig. S6.**
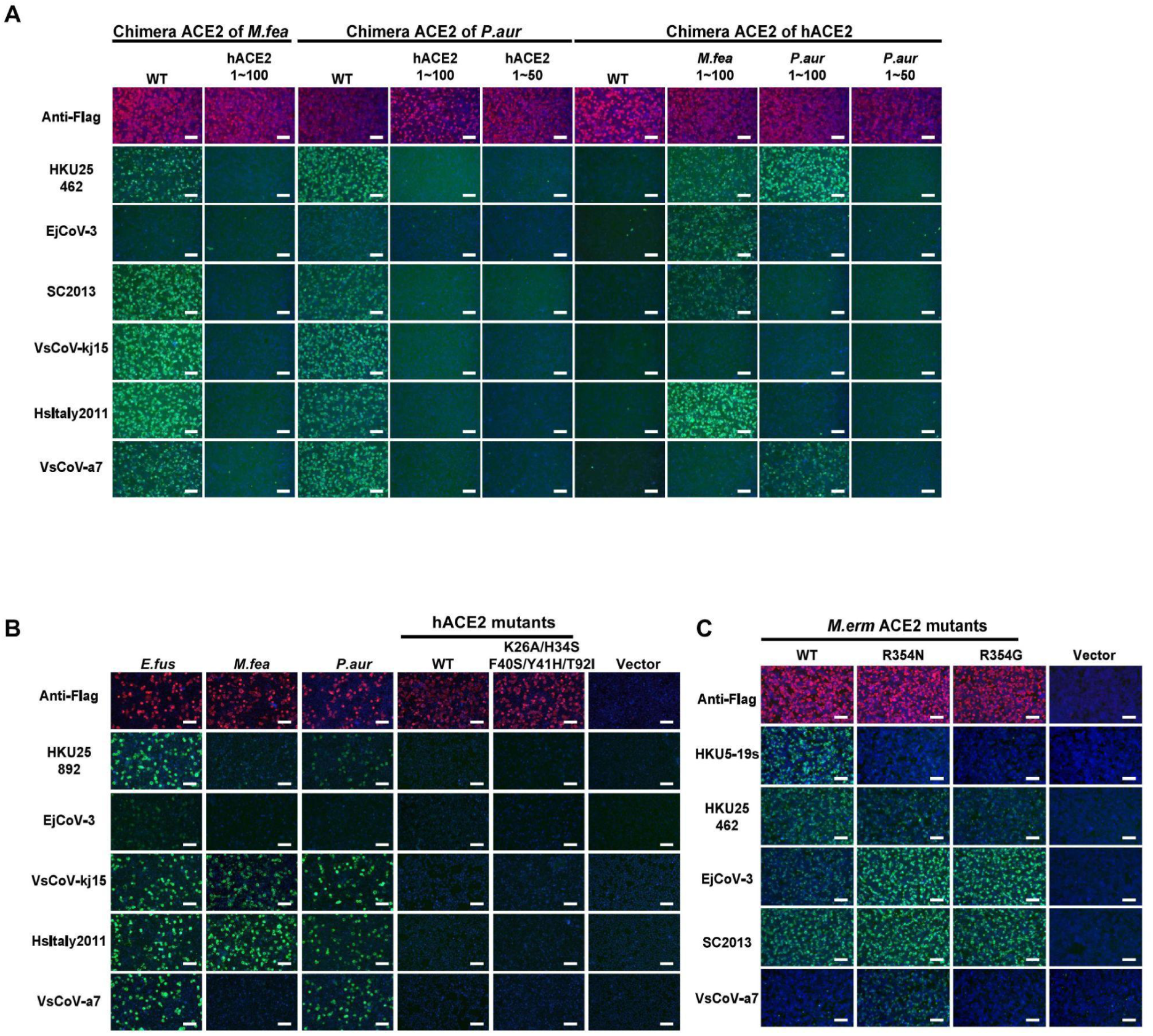
Molecular determinants of ACE2 host species tropism overlapping with HKU5. (***A***) Immunofluorescence assay analyzing HKU25r-CoV RBD binding to HEK293T cells transiently expressing ACE2 chimeras with indicated sequence swaps between hACE2 and *P.aur* ACE2 or *M.fea* ACE2. The expression levels were validated by detecting the C-terminal fused FLAG tags. (***B***) HKU25 clade viruses RBD-hFc binding to HEK293T cells transiently expressing hACE2 mutants. (***C***) HKU25 clade viruses RBD-hFc binding to HEK293T cells transiently expressing *M.erm* ACE2 mutants. Scale bars: 100 μm.

**Table S1.** Cryo-EM data collection and refinement statistics.

|  |  |  |
| --- | --- | --- |
|  | VsCoV-a7 RBD – *E.fus* ACE2<br>PDB 9N7E<br>EMD-49093 | HsItaly2011 RBD – *R.nor* ACE2<br>PDB 9N7D<br>EMD-49092 |
| **Data collection and processing** |  |  |
| Magnification | 105,000 | 105,000 |
| Voltage (kV) | 300 | 300 |
| Electron exposure (e-/Å<sup>2</sup>) | 60 | 60 |
| Defocus range (μm) | -0.2 - -3.0 | -0.2 - -3.0 |
| Pixel size (Å) | 0.843 | 0.843 |
| Symmetry imposed | C1 | C1 |
| Initial particle images (no.) | 5,485,455 | 2,124,204 |
| Final particle images (no.) | 578,871 | 831,432 |
| Map resolution (Å) | 2.5 | 2.5 |
| FSC threshold | 0.143 | 0.143 |
| **Refinement** |  |  |
| Model resolution (Å) | 2.6 | 2.6 |
| FSC threshold | 0.5 | 0.5 |
| Map sharpening *B* factor (Å<sup>2</sup>) | -86 | -77 |
| Model composition |  |  |
| Non-hydrogen atoms | 7068 | 7245 |
| Protein residues | 883 | 897 |
| Ligands | 16 | 15 |
| Water | 163 | 109 |
| *B* factors (Å<sup>2</sup>) |  |  |
| Protein | 36.7 | 30.2 |
| Ligand | 59.0 | 48.8 |
| Water | 23.8 | 20 |
| R.m.s. deviations |  |  |
| Bond lengths (Å) | 0.009 | 0.003 |
| Bond angles (°) | 0.942 | 0.593 |
| **Validation** |  |  |
| MolProbity score | 1.46 | 1.33 |
| Clashscore | 5.62 | 2.33 |
| Poor rotamers (%) | 0.97 | 2.04 |
| Ramachandran plot |  |  |
| Favored (%) | 97.14 | 97.64 |
| Allowed (%) | 2.63 | 2.25 |
| Disallowed (%) | 0.23 | 0.11 |

## Notes

### Competing Interest Statement

The authors have declared no competing interest.

## Reference

1. L. V. Tse, et al., A MERS-CoV antibody neutralizes a pre-emerging group 2c bat coronavirus. Sci Transl Med 15, eadg5567 (2023).

2. Q. Wang, et al., Bat origins of MERS-CoV supported by bat coronavirus HKU4 usage of human receptor CD26. Cell Host Microbe 16, 328–337 (2014).

3. J. Chen, et al., A bat MERS-like coronavirus circulates in pangolins and utilizes human DPP4 and host proteases for cell entry. Cell 186, 850–863.e16 (2023).

4. V. S. Raj, et al., Dipeptidyl peptidase 4 is a functional receptor for the emerging human coronavirus-EMC. Nature 495, 251–254 (2013).

5. G. Lu, et al., Molecular basis of binding between novel human coronavirus MERS-CoV and its receptor CD26. Nature 500, 227–231 (2013).

6. C.-M. Luo, et al., Discovery of Novel Bat Coronaviruses in South China That Use the Same Receptor as Middle East Respiratory Syndrome Coronavirus. J Virol 92 (2018).

7. S. K. P. Lau, et al., Receptor Usage of a Novel Bat Lineage C Betacoronavirus Reveals Evolution of Middle East Respiratory Syndrome-Related Coronavirus Spike Proteins for Human Dipeptidyl Peptidase 4 Binding. J Infect Dis 218, 197–207 (2018).

8. A. M. Zaki, S. van Boheemen, T. M. Bestebroer, A. D. M. E. Osterhaus, R. A. M. Fouchier, Isolation of a novel coronavirus from a man with pneumonia in Saudi Arabia. N Engl J Med 367, 1814–1820 (2012).

9. CSR, MERS outbreaks. World Health Organization - Regional Office for the Eastern Mediterranean. Available at: https://www.emro.who.int/health-topics/mers-cov/mers-outbreaks.html [Accessed 17 May 2024].

10. S. Mallapaty, The pathogens that could spark the next pandemic. Nature 632, 488 (2024).

11. Pathogens prioritization: a scientific framework for epidemic and pandemic research preparedness. Available at: https://www.who.int/publications/m/item/pathogens-prioritization-a-scientific-framework-for-epidemic-and-pandemic-research-preparedness [Accessed 20 January 2025].

12. Current ICTV Taxonomy Release. Available at: https://ictv.global/taxonomy [Accessed 20 January 2025].

13. H. A. Mohd, J. A. Al-Tawfiq, Z. A. Memish, Middle East Respiratory Syndrome Coronavirus (MERS-CoV) origin and animal reservoir. Virol J 13, 87 (2016).

14. J. E. Tolentino, S. Lytras, J. Ito, K. Sato, Recombination analysis on the receptor switching event of MERS-CoV and its close relatives: implications for the emergence of MERS-CoV. Virol J 21, 84 (2024).

15. V. M. Corman, et al., Rooting the phylogenetic tree of middle East respiratory syndrome coronavirus by characterization of a conspecific virus from an African bat. J Virol 88, 11297–11303 (2014).

16. M. Geldenhuys, et al., A metagenomic viral discovery approach identifies potential zoonotic and novel mammalian viruses in Neoromicia bats within South Africa. PLoS One 13, e0194527 (2018).

17. Q. Xiong, et al., Close relatives of MERS-CoV in bats use ACE2 as their functional receptors. Nature 612, 748–757 (2022).

18. Q. Xiong, et al., ACE2-using merbecoviruses: Further evidence of convergent evolution of ACE2 recognition by NeoCoV and other MERS-CoV related viruses. Cell Insight 3, 100145 (2024).

19. C.-B. Ma, et al., Multiple independent acquisitions of ACE2 usage in MERS-related coronaviruses. Cell (2025).

20. Y.-J. Park, et al., Molecular basis of convergent evolution of ACE2 receptor utilization among HKU5 coronaviruses. Cell (2025).

21. C. Ma, et al., Broad host tropism of ACE2-using MERS-related coronaviruses and determinants restricting viral recognition. Cell Discov 9, 57 (2023).

22. N. J. Catanzaro, et al., ACE2 from bats is a receptor for HKU5 coronaviruses. bioRxiv (2024).

23. J. Chen, et al., Merbecovirus HKU5 Lineage 2 Discovered in Bats Utilizes Human ACE2 as Cell Receptor. (2024). 10.2139/ssrn.4963002.

24. H. L. Wells, et al., The coronavirus recombination pathway. Cell Host Microbe 31, 874–889 (2023).

25. A. de Klerk, et al., Conserved recombination patterns across coronavirus subgenera. Virus Evol. 8, veac054 (2022).

26. Z. Zhou, Y. Qiu, X. Ge, The taxonomy, host range and pathogenicity of coronaviruses and other viruses in the Nidovirales order. Anim. Dis. 1, 5 (2021).

27. International Committee on Taxonomy of Viruses Executive Committee, The new scope of virus taxonomy: partitioning the virosphere into 15 hierarchical ranks. Nat Microbiol 5, 668–674 (2020).

28. A. Moreno, et al., Detection and full genome characterization of two beta CoV viruses related to Middle East respiratory syndrome from bats in Italy. Virol J 14, 239 (2017).

29. M. A. Wiederkehr, W. Qi, K. Schoenbaechler, C. Fraefel, J. Kubacki, Virus Diversity, Abundance, and Evolution in Three Different Bat Colonies in Switzerland. Viruses 14 (2022).

30. L. Yang, et al., MERS-related betacoronavirus in Vespertilio superans bats, China. Emerg Infect Dis 20, 1260–1262 (2014).

31. Y. Han, et al., Panoramic analysis of coronaviruses carried by representative bat species in Southern China to better understand the coronavirus sphere. Nat Commun 14, 5537 (2023).

32. S. Murakami, et al., Detection and genetic characterization of bat MERS-related coronaviruses in Japan. Transbound Emerg Dis 69, 3388–3396 (2022).

33. A. Annan, et al., Human betacoronavirus 2c EMC/2012-related viruses in bats, Ghana and Europe. Emerg Infect Dis 19, 456–459 (2013).

34. S. K. P. Lau, et al., Identification of a Novel Betacoronavirus () in Amur Hedgehogs from China. Viruses 11 (2019).

35. V. M. Corman, et al., Characterization of a novel betacoronavirus related to middle East respiratory syndrome coronavirus in European hedgehogs. J Virol 88, 717–724 (2014).

36. A. V. S. Cruz, et al., Genomic characterization and cross-species transmission potential of hedgehog coronavirus. One Health 19, 100940 (2024).

37. C. C. S. Tan, et al., Genomic screening of 16 UK native bat species through conservationist networks uncovers coronaviruses with zoonotic potential. Nat Commun 14, 3322 (2023).

38. J.-E. Park, et al., Proteolytic processing of Middle East respiratory syndrome coronavirus spikes expands virus tropism. Proc. Natl. Acad. Sci. U. S. A. 113, 12262–12267 (2016).

39. J. K. Millet, G. R. Whittaker, Host cell entry of Middle East respiratory syndrome coronavirus after two-step, furin-mediated activation of the spike protein. Proc. Natl. Acad. Sci. U. S. A. 111, 15214–15219 (2014).

40. D. Pinto, et al., Broad betacoronavirus neutralization by a stem helix-specific human antibody. Science 373, 1109–1116 (2021).

41. X. Sun, et al., Neutralization mechanism of a human antibody with pan-coronavirus reactivity including SARS-CoV-2. Nat Microbiol 7, 1063–1074 (2022).

42. S. Xia, et al., Inhibition of SARS-CoV-2 (previously 2019-nCoV) infection by a highly potent pan-coronavirus fusion inhibitor targeting its spike protein that harbors a high capacity to mediate membrane fusion. Cell Res 30, 343–355 (2020).

43. S. Xia, et al., Structural and functional basis for pan-CoV fusion inhibitors against SARS-CoV-2 and its variants with preclinical evaluation. Signal Transduct Target Ther 6, 288 (2021).

44. L. Riva, et al., Discovery of SARS-CoV-2 antiviral drugs through large-scale compound repurposing. Nature 586, 113–119 (2020).

45. M. Hoffmann, et al., SARS-CoV-2 cell entry depends on ACE2 and TMPRSS2 and is blocked by a clinically proven protease inhibitor. Cell 181, 271–280.e8 (2020).

46. Y. Du, et al., A broadly neutralizing humanized ACE2-targeting antibody against SARS-CoV-2 variants. Nat Commun 12, 5000 (2021).

47. S. Xia, et al., A pan-coronavirus fusion inhibitor targeting the HR1 domain of human coronavirus spike. Sci Adv 5, eaav4580 (2019).

48. Y. Yang, et al., Receptor usage and cell entry of bat coronavirus HKU4 provide insight into bat-to-human transmission of MERS coronavirus. Proc. Natl. Acad. Sci. U. S. A. 111, 12516–12521 (2014).

49. A. C. Walls, et al., Unexpected receptor functional mimicry elucidates activation of Coronavirus fusion. Cell 176, 1026–1039.e15 (2019).

50. P. Liu, et al., Design of customized coronavirus receptors. Nature 635, 978–986 (2024).

51. H. Guo, et al., Evolutionary Arms Race between Virus and Host Drives Genetic Diversity in Bat Severe Acute Respiratory Syndrome-Related Coronavirus Spike Genes. J Virol 94 (2020).

52. J. E. Tolentino, S. Lytras, J. Ito, E. C. Holmes, K. Sato, Recombination as an evolutionary driver of MERS-related coronavirus emergence. Lancet Infect. Dis. 24, e546 (2024).

53. J.-Y. Si, et al., Sarbecovirus RBD indels and specific residues dictating multi-species ACE2 adaptiveness. Nat Commun 15, 8869 (2024).

54. X. Yan, et al., A compendium of 8,176 bat RNA viral metagenomes reveals ecological drivers and circulation dynamics. Nat Microbiol (2025). 10.1038/s41564-024-01884-7.

55. X. Hou, et al., Using artificial intelligence to document the hidden RNA virosphere. Cell 187, 6929–6942.e16 (2024).

56. J. Wang, et al., Individual bat virome analysis reveals co-infection and spillover among bats and virus zoonotic potential. Nat Commun 14, 4079 (2023).

57. W.-T. He, et al., Virome characterization of game animals in China reveals a spectrum of emerging pathogens. Cell 185, 1117–1129.e8 (2022).

58. Y. Wang, et al., Unveiling bat-borne viruses: a comprehensive classification and analysis of virome evolution. Microbiome 12, 235 (2024).

59. J. Cui, F. Li, Z.-L. Shi, Origin and evolution of pathogenic coronaviruses. Nat Rev Microbiol 17, 181–192 (2019).

60. C. W. Tan, X. Yang, D. E. Anderson, L.-F. Wang, Bat virome research: the past, the present and the future. Curr Opin Virol 49, 68–80 (2021).

61. J. Zhao, et al., Farmed fur animals harbour viruses with zoonotic spillover potential. Nature 634, 228–233 (2024).

62. M. A. Whitt, Generation of VSV pseudotypes using recombinant ΔG-VSV for studies on virus entry, identification of entry inhibitors, and immune responses to vaccines. J. Virol. Methods 169, 365–374 (2010).

63. S. Biacchesi, et al., Rapid human metapneumovirus microneutralization assay based on green fluorescent protein expression. J. Virol. Methods 128, 192–197 (2005).

64. J. Nie, et al., Quantification of SARS-CoV-2 neutralizing antibody by a pseudotyped virus-based assay. Nat. Protoc. 15, 3699–3715 (2020).

65. H. Yan, et al., ACE2 receptor usage reveals variation in susceptibility to SARS-CoV and SARS-CoV-2 infection among bat species. *Nat*. Ecol. Evol. 5, 600–608 (2021).

66. Y. Liu, et al., Functional and genetic analysis of viral receptor ACE2 orthologs reveals a broad potential host range of SARS-CoV-2. Proc. Natl. Acad. Sci. U. S. A. 118 (2021).

67. C. Schwegmann-Weßels, et al., Comparison of vesicular stomatitis virus pseudotyped with the S proteins from a porcine and a human coronavirus. J Gen Virol 90, 1724–1729 (2009).

68. C. J. Russo, L. A. Passmore, Electron microscopy: Ultrastable gold substrates for electron cryomicroscopy. Science 346, 1377–1380 (2014).

69. C. Suloway, et al., Automated molecular microscopy: the new Leginon system. J. Struct. Biol. 151, 41–60 (2005).

70. D. Tegunov, P. Cramer, Real-time cryo-electron microscopy data preprocessing with Warp. Nat. Methods 16, 1146–1152 (2019).

71. A. Punjani, J. L. Rubinstein, D. J. Fleet, M. A. Brubaker, cryoSPARC: algorithms for rapid unsupervised cryo-EM structure determination. Nat. Methods 14, 290–296 (2017).

72. T. Bepler, et al., Positive-unlabeled convolutional neural networks for particle picking in cryo-electron micrographs. Nat. Methods 16, 1153–1160 (2019).

73. A. Punjani, H. Zhang, D. J. Fleet, Non-uniform refinement: adaptive regularization improves single-particle cryo-EM reconstruction. Nat. Methods 17, 1214–1221 (2020).

74. J. Zivanov, et al., New tools for automated high-resolution cryo-EM structure determination in RELION-3. Elife 7 (2018).

75. P. B. Rosenthal, R. Henderson, Optimal determination of particle orientation, absolute hand, and contrast loss in single-particle electron cryomicroscopy. J. Mol. Biol. 333, 721–745 (2003).

76. S. Chen, et al., High-resolution noise substitution to measure overfitting and validate resolution in 3D structure determination by single particle electron cryomicroscopy. Ultramicroscopy 135, 24–35 (2013).

77. E. F. Pettersen, et al., UCSF Chimera--a visualization system for exploratory research and analysis. J. Comput. Chem. 25, 1605–1612 (2004).

78. P. Emsley, B. Lohkamp, W. G. Scott, K. Cowtan, Features and development of coot. Acta Crystallogr. D Biol. Crystallogr. 66, 486–501 (2010).

79. J. Abramson, et al., Accurate structure prediction of biomolecular interactions with AlphaFold 3. Nature 630, 493–500 (2024).

80. D. Liebschner, et al., Macromolecular structure determination using X-rays, neutrons and electrons: recent developments in Phenix. Acta Crystallogr. D Struct. Biol. 75, 861–877 (2019).

81. V. B. Chen, et al., MolProbity: all-atom structure validation for macromolecular crystallography. Acta Crystallogr. D Biol. Crystallogr. 66, 12–21 (2010).

82. B. A. Barad, et al., EMRinger: side chain-directed model and map validation for 3D cryo-electron microscopy. Nat. Methods 12, 943–946 (2015).

83. J. Agirre, et al., Privateer: software for the conformational validation of carbohydrate structures. Nat. Struct. Mol. Biol. 22, 833–834 (2015).

